# A shuttle-vector system allows heterologous gene expression in the thermophilic methanogen *Methanothermobacter thermautotrophicus* ΔH

**DOI:** 10.1101/2021.04.20.440605

**Authors:** Christian Fink, Sebastian Beblawy, Andreas M. Enkerlin, Lucas Mühling, Largus T. Angenent, Bastian Molitor

## Abstract

Thermophilic *Methanothermobacter* spp. are used as model microbes to study the physiology and biochemistry of the conversion of hydrogen and carbon dioxide into methane (*i.e*., hydrogenotrophic methanogenesis), because of their short doubling times and robust growth with high growth yields. Yet, a genetic system for these model microbes was missing despite intense work for four decades. Here, we report the establishment of tools for genetic modification of *M. thermautotrophicus*. We developed the modular *Methanothermobacter* vector system, which provided shuttle-vector plasmids (pMVS) with exchangeable selectable markers and replicons for both *Escherichia coli* and *M. thermautotrophicus*. For *M. thermautotrophicus*, a thermostable neomycin-resistance cassette served as the selectable marker for positive selection with neomycin, and the cryptic plasmid pME2001 from *Methanothermobacter marburgensis* served as the replicon. The pMVS-plasmid DNA was transferred from *E. coli* into *M. thermautotrophicus via* interdomain conjugation. After the successful validation of DNA transfer and positive selection in *M. thermautotrophicus*, we demonstrated heterologous gene expression of a thermostable β-galactosidase-encoding gene (*bgaB*) from *Geobacillus stearothermophilus* under the expression control of four distinct synthetic and native promoters. In quantitative *in-vitro* enzyme activity assays, we found significantly different β-galactosidase activity with these distinct promoters. With a formate dehydrogenase operon-encoding shuttle vector, we allowed growth of *M. thermautotrophicus* on formate as the sole growth substrate, while this was not possible for the empty vector control. These genetic tools provide the basis to investigate hypotheses from four decades of research on the physiology and biochemistry of *Methanothermobacter* spp. on a genetic level.

**Significance Statement:** The world economies are facing permanently increasing energy demands. At the same time, carbon emissions from fossil sources need to be circumvented to minimize harmful effects from climate change. The power-to-gas platform is utilized to store renewable electric power and decarbonize the natural gas grid. The microbe *Methanothermobacter thermautotrophicus* is already applied as the industrial biocatalyst for the biological methanation step in large-scale power-to-gas processes. To improve the biocatalyst in a targeted fashion, genetic engineering is required. With our shuttle-vector system for heterologous gene expression in *M. thermautotrophicus*, we set the cornerstone to engineer the microbe for optimized methane production, but also for production of high-value platform chemicals in power-to-x processes.

## Introduction

Methanogenesis is the biological production of methane, which is catalyzed by methane-producing archaea (methanogens). Hydrogenotrophic methanogens grow with hydrogen (electron donor) and carbon dioxide (electron acceptor and carbon source) as the substrates (1). For decades, the Methanobacteriales species *Methanothermobacter thermautotrophicus* and *Methanothermobacter marburgensis* have been studied as model microbes for the biochemistry of the hydrogenotrophic methanogenic metabolism, and deep insights into their energy and carbon metabolism have been acquired (1–3). For example, comparative genome analyses of *M. thermautotrophicus* and *M. marburgensis* revealed the genes that are most likely required for hydrogenotrophic methanogenesis (4), and a plethora of studies unraveled the mechanism of key enzymes such as the methyl-coenzyme M reductase (reviewed in (5)). Furthermore, the pseudomurein-containing cell wall, which is specific to Methanobacteriales, has attracted research on *Methanothermobacter* spp. (6). This long-lasting interest in the biochemistry and physiology of these model microbes led to extensive attempts to establish genetic tools for *Methanothermobacter* spp. in the past (7–9). Successes were reported in *M. marburgensis* for reversing of spontaneous amino acid-auxotrophic phenotypes *via* generalized transduction from the wild-type strain with the virus ΨM2 (10), or *via* the uptake of free DNA (*i.e*., natural competence) with high-molecular weight genomic DNA (11–13). In addition, 5-fluoro-uracil-resistant phenotypes were reported to be conferred with high-molecular weight genomic DNA from a spontaneous-resistant strain *via* natural competence in *M. marburgensis* with a higher frequency than the rate for the occurrence of spontaneous resistance (11). Other spontaneous antibiotic-resistant *Methanothermobacter* strains were investigated as potential sources for genes that can be used as selectable markers, including pseudomonic acid- (12) and neomycin- (14) resistant strains. Furthermore, several potential *Escherichia coli-Methanothermobacter* shuttle vectors have been constructed and replicated in *E. coli*, which were based on the cryptic plasmid (*i.e*., no physiological function has been assigned to the plasmid) pME2001 from *M. marburgensis* (7). Nevertheless, until now none of these approaches was translated into a reliable genetic system for *Methanothermobacter* spp.

In contrast, genetic systems for mesophilic methanogens have been well established for *Methanococcus* spp. and *Methanosarcina* spp. Sophisticated genetic tools for these microbes have been implemented, such as allelic exchange methods and counter-selection markers to generate gene deletion strains, and recently CRISPR/Cas9-based genome engineering (15–17). The transfer of DNA into the cells for these systems has been achieved with varying efficiencies *via*: **1)** natural competence (18); **2)** polyethylene glycol-mediated transformation of protoplasts (16); **3)** liposome-mediated transformation of protoplasts (19); **4)** electroporation (20); and **5)** conjugation (21). As antibiotic-selectable markers, a puromycin-selectable marker (19), different kanamycin/neomycin-selectable markers (22), a mupirocin-selectable marker (12), and a nourseothricin-selectable marker (23) have been implemented. These selectable markers are typically placed under the control of the constitutive promoter sequence, which is located upstream of the methyl-coenzyme M reductase (*mcr*) operon from *Methanococcus voltae* (P_*mcrB*(*M.v*.)_) (16), for both *Methanococcus* spp. and *Methanosarcina* spp. An exception is the nourseothricin marker, which has been placed under the control of a modified version of the *mcr* promoter from *Methanosarcina barkeri* (P_*mcrB*-tetO1(*M.b*.)_) (24). This selectable marker has been only tested for *Methanosarcina* spp. so far (23). Furthermore, shuttle vectors that can be replicated in *E. coli* and methanogens have been established based on the cryptic plasmid pURB500 for *M. maripaludis* (25), and based on the cryptic plasmid pC2A for *Methanosarcina* spp. (19).

In addition to the mesophilic systems, genetic tools for the thermophilic methanogens *Methanocaldococcus jannaschii* and *Methanoculleus thermophilus* have recently been described (18, 26). DNA transfer was achieved *via* heat shock-mediated transformation (*M. jannaschii*) or natural competence (*M. thermophilus*). Positive selection with antibiotics was accomplished for *M. jannaschii* with a simvastatin/mevilonin-selectable marker under the control of the native constitutive promoter of an S-Layer protein-encoding gene (P_Sla(*M.j*.)_) (26), and for *M. thermophilus* with a thermostability-adapted neomycin-selectable marker (27) under the control of the promoter of the methyl-tetrahydromethanopterin:coenzyme M methyl-transferase (*mtr*) operon from *M. jannaschii* (P_*mtr*(*M.j*.)_) (18). Both systems rely on the integration of DNA into the genome by homologous recombination, and replicating plasmid vectors for the thermophilic methanogens have not been described.

Despite the fact that no genetic system for *M. thermautotrophicus* and *M. marburgensis* was available so far, wild-type cultures of those microbes have been implemented as biocatalysts in the power-to-gas platform (28, 29), because high cell densities and high methane production rates can be achieved (30, 31). For example, Electrochaea GmbH (Planegg, Germany) uses a proprietary (non-genetically modified) *M. thermautotrophicus* wild-type strain (32). In the power-to-gas platform, hydrogen from the electrolysis of water with surplus electric power from renewable resources is combined with carbon dioxide, and these gases are converted into methane (28). This methane (*i.e*., renewable natural gas) can be introduced into the natural gas grid with large storage capacities to substitute fossil natural gas.

Genetic engineering will be necessary to further optimize the biocatalysts for power-to-gas processes in a targeted fashion. With genetic engineering, the metabolism of the biocatalysts can, for example, be improved to maximize methane-production rates, or be amended for an expanded substrate or product spectrum, which would allow to convert the power-to-gas into a broader power-to-chemicals (*i.e*., power-to-x) platform (29). Spurred by the recurring interest in *Methanothermobacter* spp. as model microbes for methanogenic biochemistry and physiology with a long history, and as biotechnologically relevant microbes in power-to-gas processes, we set out to utilize modern molecular biology tools and the knowledge on genetic tools for other methanogens and thermophilic microbes to develop a genetic system for *M. thermautotrophicus* ΔH. Here, we report the successful implementation of genetic tools for *M. thermautotrophicus* ΔH, including: **1)** a shuttle vector that replicates in *E. coli* and *M. thermautotrophicus* ΔH; **2)** an interdomain conjugation protocol for conjugational DNA transfer from *E. coli* S17-1 to *M. thermautotrophicus* ΔH; and **3)** a neomycin-selectable marker. We further demonstrated the functionality of these genetic tools by implementing a thermostable β-galactosidase as a reporter for genetic studies. Finally, we amended the metabolism of *M. thermautotrophicus* ΔH to allow growth on formate as an alternative substrate to demonstrate the potential of our genetic tools for biotechnological applications.

## Results

### Clonal populations of *M. thermautotrophicus* ΔH can be obtained on solidified media plates as individual colonies with high plating efficiencies

The first requirement to allow genetic work with any given microbe is the capability to isolate clonal populations. This is typically achieved by plating microbial cultures on solidified media plates and by selecting individual colonies. Therefore, we first reproduced the high plating efficiencies that have been reported in the literature for *Methanothermobacter* spp. (33). We investigated three common plating techniques (spot-, spread-, and pour-plating), and compared factors that influenced plating efficiency (*i.e*., individual colonies per cell-count of plated microbial cells, ***Materials and Methods***). We achieved dense growth with spot-plating, but individual colonies were barely distinguishable with this plating technique, while we achieved plating efficiencies between 1-5% with spread-plating, and 50% and higher with pour-plating (***SI Appendix*, *Results S1*, *Figure S1***).

### *M. thermautotrophicus* ΔH is sensitive to antibiotics commonly used in methanogen genetic systems

To find a suitable selection pressure for the positive selection of genetically modified cells, we analyzed several antibiotics such as simvastatin and neomycin (***SI Appendix*, Results S2**). For both antibiotics, thermostable selectable markers are available, which have been successfully used in thermophilic non-methanogenic microbes, such as *Pyrococcus furiosus*, *Thermococcus kodakarensis* (simvastatin) (34, 35), as well as *Thermoanaerobacter* spp. (neomycin) (36), but recently also in the thermophilic methanogens *M. jannaschii* (simvastatin) and *M. thermophilus* (neomycin) (18, 26). Both simvastatin and neomycin efficiently inhibited growth of *M. thermautotrophicus* ΔH cells in liquid culture at concentrations of 21.5 μg/mL and 250 μg/mL, respectively (***SI Appendix*, Results S2, Figure S2 and S3**). While these antibiotics inhibited growth of *M. thermautotrophicus* ΔH, we observed the appearance of spontaneous-resistant *M. thermautotrophicus* ΔH cells for both antibiotics (***SI Appendix*, Results S2, Figure S2 and S3**). The incubation period for the appearance of spontaneous-resistant *M. thermautotrophicus* ΔH cells differed for each antibiotic, when compared to the incubation period for non-selective growth of wild-type cells (16-24 h). We observed inhibition of growth for at least 48 h with simvastatin, and 60 h with neomycin at the concentrations indicated above (***SI Appendix*, Results S2**). Because we found that neomycin inhibits growth of *M. thermautotrophicus* ΔH for a longer incubation period compared to simvastatin, we decided to focus on neomycin as the selection pressure to develop a first selectable marker for *M. thermautotrophicus* ΔH.

### A modular plasmid design enables the fast generation of shuttle vectors to test genetic elements for functionality in *M. thermautotrophicus* ΔH

Before we focused on the neomycin-selectable marker, we had constructed a subset of plasmids that would allow to test various approaches for the transfer of DNA into *M. thermautotrophicus* ΔH and for positive selection (***Materials and Methods***). To ease exchangeability of genetic elements in shuttle vectors, and to allow fast adaptation to new findings, we decided for a modular plasmid design from the beginning. Inspired by the pSEVA system for Gram-negative bacteria (37), and the pMTL80000 system for Clostridia (38), we established the *Methanothermobacter* vector system (pMVS) design.

The pMVS design consists of five modules, which are separated by rare eight base-pair recognition sequences for the restriction enzymes *Pme*I, *Asi*SI, *Fse*I, *Asc*I, and *Pac*I (**Figure 1**). To follow the pMVS design, these rare restriction enzyme-recognition sequences need to stay unique to grant exchangeability of the modules by restriction/ligation cloning. The five modules are (restriction enzyme boundaries are given in parenthesis): **1)** the replicon for *E. coli* (*Asi*SI, *Pme*I); **2)** the selectable marker for *E. coli* (*Pme*I, *Fse*I); **3)** the replicon for *M. thermautotrophicus* ΔH (*Asi*SI, *Pac*I); **4)** the selectable marker for *M. thermautotrophicus* ΔH (*Asc*I, *Fse*I); and **5)** an application module that can be used to include any genetic cargo such as a reporter gene or another gene of interest (*Pac*I, *Asc*I) (**Figure 1; *SI Appendix*, Results S3**).

**Figure 1.**
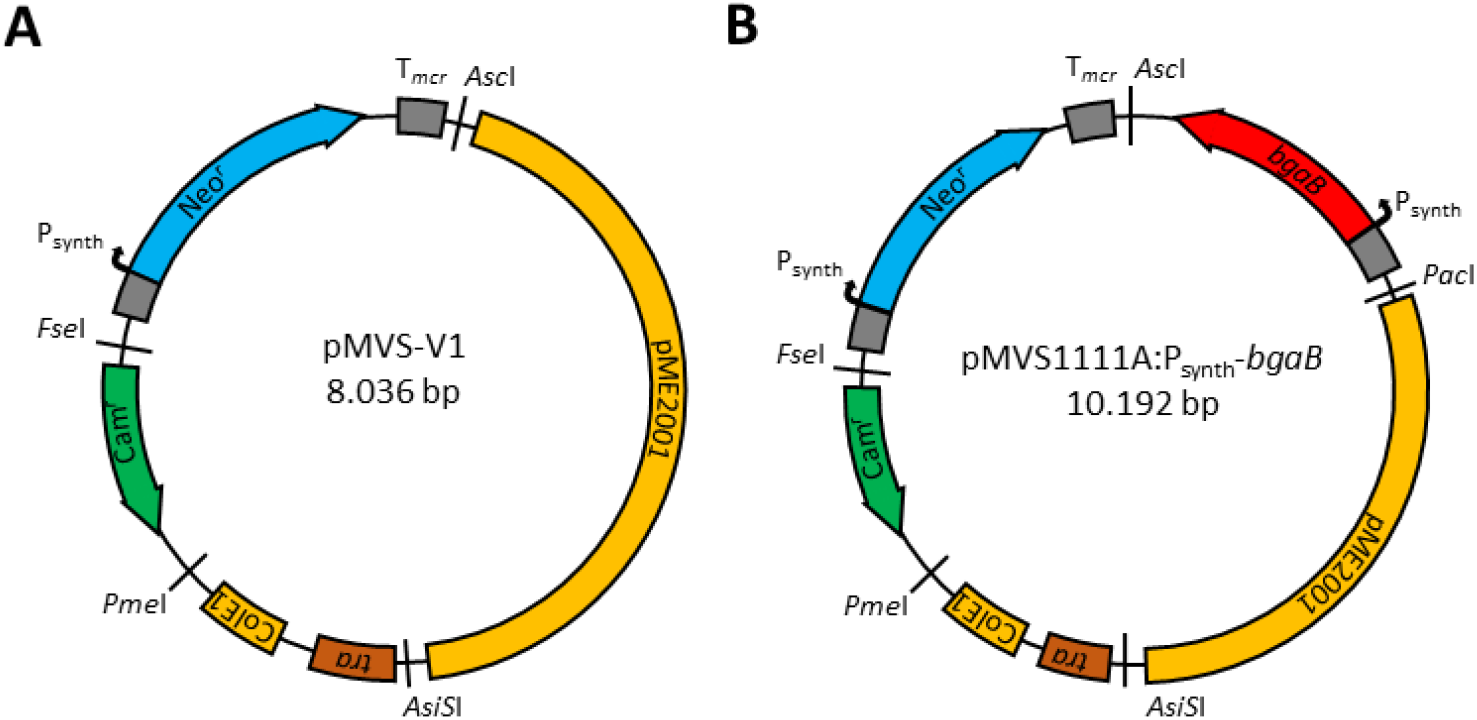
Plasmid maps of the *Methanothermobacter* vector system (pMVS). **A)** pMVS-V1 consists of four modules, which are intersected by the eight base-pair restriction enzyme recognition sites *Asc*I, *Fse*I, *Pme*I, and *Asi*SI. The four modules are the replicon for *E. coli* (ColE1, *tra*), the selectable marker for *E. coli* (Cam^r^), the replicon for *M. thermautotrophicus* ΔH (pME2001), and the selectable marker for *M. thermautotrophicus* ΔH (Neo^r^). **B)** pMVS1111A:P_synth_-*bgaB* consists of five modules, which are intersected by the eight base-pair restriction enzyme recognition sites *Asc*I, *Fse*I, *Pme*I, *Asi*SI, and *Pac*I. The five modules are as in pMVS-V1 with following differences: the replicon for *M. thermautotrophicus* ΔH is flanked by *AsiS*I and *Pac*I, and the shuttle vector contains the *bgaB* gene in the application module, which is flanked by *Pac*I and *Asc*I. P_synth_, synthetic promoter sequence, which is based on P_*hmtB*_; T_*mcr*_, terminator sequence from the *mcr* operon of *M. voltae*. The nomenclature for the pMVS design is realized by adding a four-digit code after pMVS for the definition of the first four modules, which defines the plasmid backbone with basic functions for replication and selection in *E. coli* (module 1 and 2) and *M. thermautotrophicus* ΔH (module 3 and 4). Additional large capital letters can be amended to each digit to define differences, such as varying promoter sequences, in a given module. For the fifth (application) module, a descriptive name is added after the four-digit code, which allows to stay as flexible as possible with the used genetic cargo (without a limitation to nine digits), while staying within the pMVS design boundaries.

Based on the first shuttle vector (pMVS-V1) that led to successful DNA transfer and selection protocols, as described in the next section, we defined pMVS-V1 as our archetype shuttle vector (**Figure 1A**), with a combination of: **1)** the ColE1-derived replicon for *E. coli* in combination with the conjugational transfer function (*tra*-region) from RK2 (38); **2)** the chloramphenicol-selectable marker (Cam^r^) for *E. coli* (38); **3)** the entire cryptic plasmid pME2001 from *M. marburgensis* as the replicon for *M. thermautotrophicus* ΔH (9); and **4)** the thermostable neomycin-selectable marker (Neo^r^) (27) for *M. thermautotrophicus* ΔH under the control of the P_synth_ promoter sequence (35), but without an application module and a *Pac*I-recognition sequence (**Figure 1A**). After we had demonstrated the functionality of pMVS-V1, our first complete shuttle vector, pMVS1111A-P_synth_-*bgaB*, was then constructed based on this archetype shuttle vector. pMVS1111A-P_synth_-*bgaB* contains the β-galactosidase-encoding gene *bgaB* (see below) and an additional *Pac*I site, which completes the application module (**Figure 1B; *Materials and Methods; SI Appendix*, Results S3**).

### DNA transfer into *M. thermautotrophicus* ΔH is possible *via* interdomain conjugation with *E. coli* S17-1

Following our broad approach, we investigated several published protocols (and modified versions) to transfer DNA into *M. thermautotrophicus* ΔH by using our various plasmids and shuttle vectors (***Material and Methods***). These DNA-transfer protocols included: **1)** natural competence protocols (11, 26); **2)** chemically/physically-induced transformation protocols, such as with elevated calcium and magnesium concentrations in the mineral medium, or with low temperature incubation of pre-cultures to induce stress conditions; **3)** an electroporation protocol (39); and **4)** an interdomain conjugation protocol with *E. coli* (21). Most attempts did not result in cells with the expected selectable phenotype, and if so, could not be linked to the respective anticipated genotype, and appeared to be spontaneous-resistant cells. The protocol that finally led to a successful DNA transfer into *M. thermautotrophicus* ΔH was an interdomain conjugation protocol with *E. coli* S17-1 (**Figure 2; *Materials and Methods***), which was a modified version of the protocol for conjugational DNA transfer into *M. maripaludis* (21).

**Figure 2.**
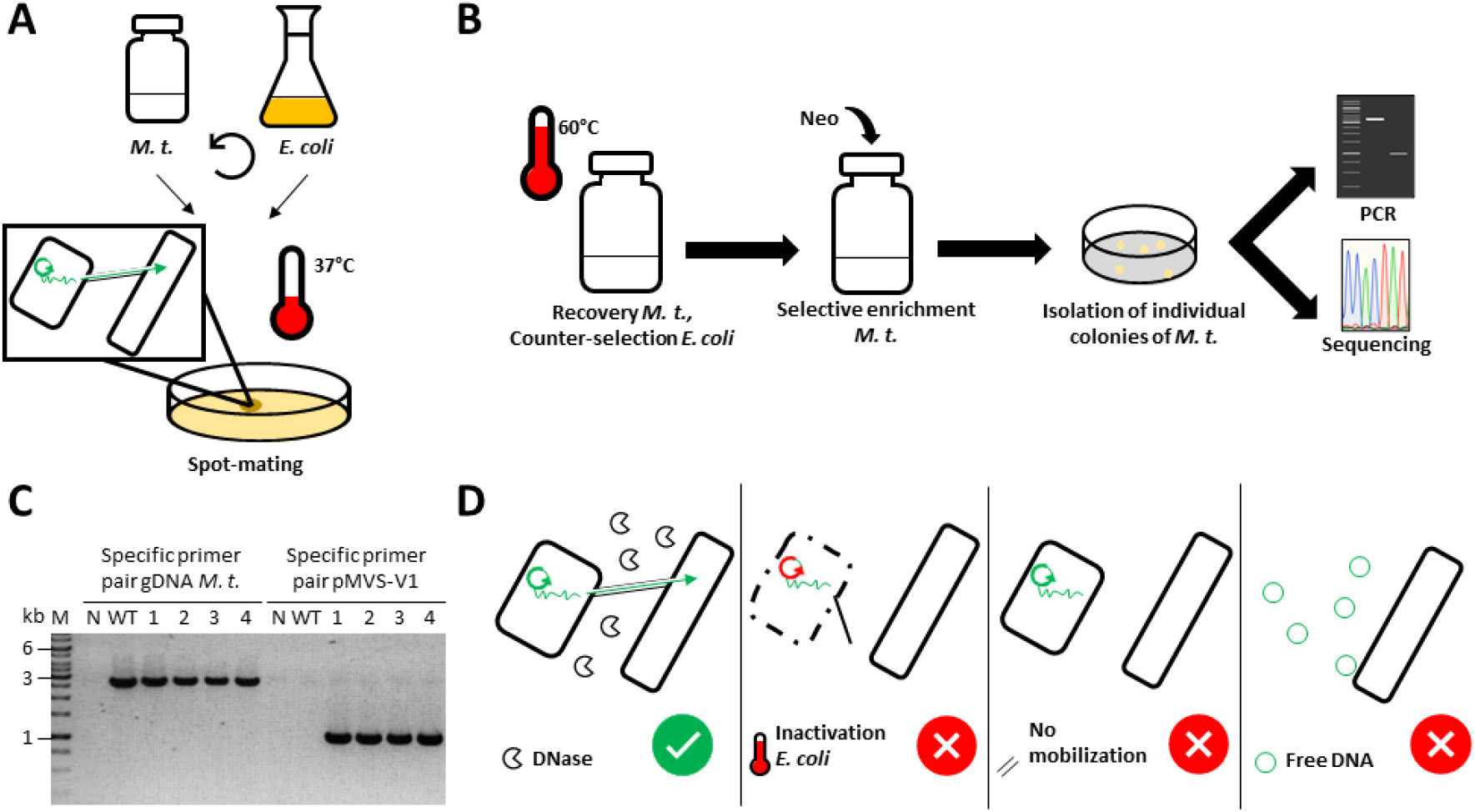
Schematic depiction and analysis of interdomain conjugation between *E. coli* S17-1 and *M. thermautotrophicus* ΔH. **A)** Wild-type *M. thermautotrophicus* ΔH (*M. t*.) and the shuttle vector-carrying *E. coli* were harvested by centrifugation, mixed, and spotted on solidified media plates that support growth of both microbes. During the spot-mating step at 37°C, the DNA-transfer process *via* conjugation takes place (small scheme). **B)** The process to isolate and identify individual colonies of genetically modified *M. thermautotrophicus* ΔH in the standard protocol. After the spot-mating step, *M. thermautotrophicus* ΔH cells were recovered in non-selective liquid mineral medium at 60°C, and afterwards, transconjugants were enriched in neomycin (Neo)-containing selective liquid mineral medium at 60°C. Individual colonies were obtained from plating the enrichment culture. Those colonies were analyzed by PCR and Sanger sequencing. **C)** PCR analysis of four respective transconjugants (**1-4**) with primer combinations specific for the shuttle vector pMVS-V1 replicon (1-kilobase fragment) and for genomic DNA of *M. thermautotrophicus* ΔH (2.8-kilobase fragment). **N**, water negative control; **WT**, control with wild-type *M. thermautotrophicus* ΔH; **M**, GeneRuler 1 kb DNA Ladder (Thermo Scientific, Waltham MA, USA). **D)** Experimental conditions for the confirmation of conjugation as the mechanism for DNA transfer were (from left to right): DNAse-I treatment, heat inactivation of *E. coli* S17-1, conjugation with non-conjugative *E. coli* NEB stable, and addition of free plasmid DNA directly to *M. thermautotrophicus* ΔH cell culture.

To achieve a successful conjugational DNA transfer of the archetype shuttle vector pMVS-V1 from *E. coli* S17-1 (donor) to *M. thermautotrophicus* ΔH (recipient), we increased the recipient cell concentration to ~1.6·10^9^ cells from a culture in the early stationary growth phase, which is a 5-fold higher cell concentration compared to Dodsworth *et al*. (2010). Furthermore, we used a spot-mating procedure to allow close physical contact between donor and recipient cells during an incubation period at 37°C on solidified media plates, which supported metabolic activity of *E. coli* S17-1 (**Figure 2A**), in contrast to direct spread-plating as in Dodsworth *et al*. (2010). After resuspending the spot-mated cells (donor + recipient) from the solidified media plate, we recovered *M. thermautotrophicus* ΔH in liquid mineral medium without any complex media additions (no organic carbon source) and without antibiotics additions at 60°C under a hydrogen/carbon dioxide atmosphere (**Figure 2B; *Materials and Methods***). These incubation conditions decreased the viability of *E. coli* S17-1 to a very minimum, and therefore no counter-selection with antibiotics against donor cells was required. After a short incubation period (3-4 h) to recover *M. thermautotrophicus* ΔH under non-selective conditions, the cells were transferred to liquid medium, which contained neomycin for a selective-enrichment step. Importantly, the required incubation period for the cells to grow in this step was key for a successful identification of transconjugants (*i.e*., recipient cells that received the shuttle vector). Neomycin at a concentration of 250 μg/mL inhibits growth of *M. thermautotrophicus* ΔH for at least 60 h in liquid medium (***SI Appendix*, Results S2**). Therefore, when the cells did not grow within less than 60 h (typically growth appeared after 24-48 h in a successful experiment) in the selective-enrichment step, the conjugation experiment was regarded as unsuccessful, because the number of spontaneous-resistant cells is considerably higher compared to potential transconjugants, which renders screening essentially impossible.

### The shuttle-vector DNA confers the observed antibiotic-resistant phenotype and is maintained in *M. thermautotrophicus* ΔH with high segregational stability

After we found selective growth of putative transconjugant cells, we confirmed the successful DNA transfer into *M. thermautotrophicus* ΔH *via* two site-specific PCR reactions, which amplified a fragment of the pMVS-V1 shuttle vector, and a fragment of the *M. thermautotrophicus* ΔH genomic DNA, respectively, with liquid cultures derived from individual colonies (**Figure 2C**). In addition, we extracted plasmid DNA from several independent *M. thermautotrophicus* ΔH transconjugant cultures, transformed *E. coli* with this plasmid-DNA extract, re-extracted the plasmid DNA again from *E. coli*, and finally, performed restriction-enzyme digestions and Sanger sequencing to confirm the plasmid-DNA integrity and sequence, without finding any differences when compared to the original pMVS-V1 shuttle vector (***SI Appendix*, Results S5, Figure S4**). With different shuttle vectors in independent experiments, we achieved reliable DNA transfer into *M. thermautotrophicus* ΔH with our standard protocol, which includes a selective-enrichment step (**Figure 2B**). To determine the conjugation frequencies, we performed experiments without the selective-enrichment step, but with a prolonged non-selective-recovery step, and we achieved conjugation frequencies of approximately 4·10^-9^ to 6·10^-6^ transconjugants per initial recipients, while experimental variations considerably influenced these numbers (***SI Appendix*, Results S6, Table S1**). Once the plasmid DNA was transferred into *M. thermautotrophicus* ΔH, it was maintained with high segregational stability over many cell divisions in an experiment under non-selective growth conditions, and we did not observe loss of pMVS-V1 (***SI Appendix*, Results S7, Figure S5**).

### Free plasmid DNA is not resulting in DNA transfer into *M. thermautotrophicus* ΔH

By having demonstrated DNA transfer into *M. thermautotrophicus* ΔH, we further analyzed whether this transfer was indeed depending on conjugational DNA transfer from *E. coli* or whether it was rather by uptake of free DNA under the utilized cultivation conditions during the conjugation protocol (***Materials and Methods***). *E. coli* S17-1 donor cells contain a large amount of pMVS-V1, because of the high-copy number ColE1 replicon, which might be released into the liquid medium from lysing cells. Therefore, we conducted control experiments with free pMVS-V1 plasmid DNA, heat-inactivated *E. coli* S17-1 cells, and non-conjugative *E. coli* NEB stable cells that carry pMVS-V1 (**Figure 2D**). None of these experiments resulted in DNA transfer into *M. thermautotrophicus* ΔH (**Figure 2D; *SI Appendix*, Results S8, Figure S6**). In contrast, DNAse-I treatment of the *E. coli* S17-1 donor cells did not negatively influence the success of a conjugational DNA transfer into *M. thermautotrophicus* ΔH (**Figure 2D; *SI Appendix*, Results S8, Figure S6**). Thus, we concluded that DNA transfer occurs due to conjugational mobilization activity from *E. coli* S17-1 to *M. thermautotrophicus* ΔH.

### A thermostable ß-galactosidase (BgaB) from *Geobacillus stearothermophilus* is a functional reporter to investigate promoter sequences in *M. thermautotrophicus* ΔH

With a DNA-transfer protocol and a functional shuttle vector at hands, we proceeded with adding a genetic cargo (*i.e*., gene of interest) to the application module of the archetype pMVS-V1 shuttle vector. To enable the analysis of the effects from different promoter sequences on gene expression in *M. thermautotrophicus* ΔH, we decided to implement a reporter gene as our first gene of interest. We chose the *bgaB* gene from *Geobacillus stearothermophilus*, which encodes a thermostable β-galactosidase (40). We placed a codon-optimized version of the *bgaB* gene under the control of the non-native P_synth_ promoter (***Materials and Methods***). We transferred the resulting shuttle vector pMVS1111A:P_synth_-*bgaB* (**Figure 1B**) into *M. thermautotrophicus* ΔH *via* conjugation. In a qualitative preliminary experiment with cell lysate from pMVS1111A:P_synth_-*bgaB*-carrying *M. thermautotrophicus* ΔH cells (and pMVS-V1-carrying cells as an empty vector negative control), we found that, indeed, the β-galactosidase BgaB is produced in *M. thermautotrophicus* ΔH and results in a color reaction in an enzyme assay with 3,4-cyclohexenoesculetin-β-D-galactopyranoside (S-Gal) only in the presence of the *bgaB* gene (**Figure 3B; *Materials and Methods***).

**Figure 3.**
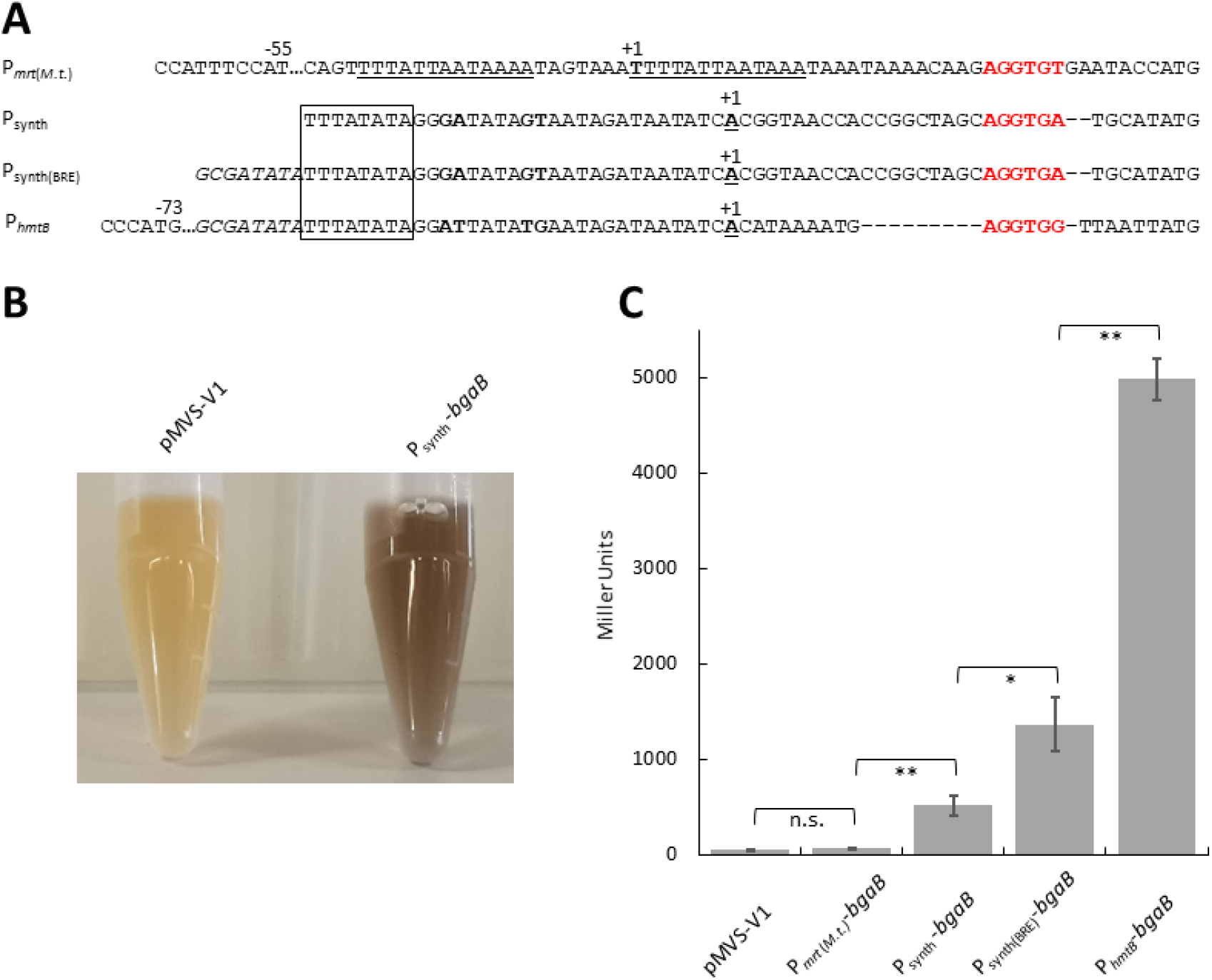
Enzyme activity assays with *M. thermautotrophicus* ΔH strains that carry a thermostable β-galactosidase (BgaB)-encoding gene under the control of four distinct promoter sequences. **A)** Sequence alignment of distinct putative promoter sequences that we analyzed for activity to drive the expression of a thermostable β-galactosidase (*bgaB*) gene. Sequence repeats in P_*mrt*(*M.t*.)_ are underlined. The transcription start site is indicated by “+1”, highlighted in bold, and underlined. TATA-box sequences of P_synth_, P_synth(BRE)_, and P_*hmtB*_ are surrounded by a box. BRE sequences are highlighted in italics and ribosome-binding sites in red. Dashes are used as spacers, while dots indicate additional base pairs, which are left out here for visualization. Differences between P_synth_, P_synth(BRE)_, and P_*hmtB*_ between the TATA-box sequence and transcription start site are highlighted in bold. **B)** Qualitative analysis of BgaB activity with S-Gal as chromogenic substance in an *in-vitro* assay with cell lysate of empty vector-carrying *M. thermautotrophicus* ΔH (pMVS-V1) or pMVS1111A:P_synth_-*bgaB*-carrying *M. thermautotrophicus* ΔH (P_synth_-*bgaB*) cells. **C)** Quantitative analysis of BgaB activity with ONPG as chromogenic substance in an *in-vitro* assay with cell lysate of *M. thermautotrophicus* ΔH strains that carry plasmids with the *bgaB* gene under the control of the four distinct promoters (pMVS-V1, empty vector control; P_*mrt*(*M.t*.)_-*bgaB*, pMVS1111A:P_*mrt*(*M.t*.)_-*bgaB*; P_synth_-*bgaB*, pMVS1111A:P_synth_-*bgaB*; P_synth(BRE)_-*bgaB*, pMVS1111A: P_synth(BRE)_-*bgaB*; P_*hmtB*_-*bgaB* pMVS1111A:P_*hmtB*_-*bgaB*). Average (*N*=3) with error bars indicating standard deviation. Significance was tested with Student’s t-test (two-tailed): *, significant difference (P<0.05); **, highly significant difference (P<0.01); n.s., no significant difference (P>0.05).

This result sparked us to establish a quantitative β-galactosidase enzyme activity assay with o-nitrophenyl-β-D-galactopyranoside (ONPG) as the chromogenic substrate for the β-galactosidase (***SI Appendix*, Results S9**), which allowed us to investigate promoter sequences for their relative *in-vivo* effect on gene expression in *M. thermautotrophicus* ΔH during a growth experiment (***SI Appendix*, Figure S7**). Overall, we selected four distinct promoter sequences (P_synth_, P_synth(BRE)_, P_*hmtB*_, and P_*mrt*(*M.t*.)_) based on our previous results and the peer-reviewed literature, and compared the effects of the promoters on gene expression with the established enzyme assay (**Figure 3; *SI Appendix*, Results S9**). With our optimized quantitative β-galactosidase enzyme activity assay, we found that the P_synth_ promoter, without a transcription factor B recognition element (BRE) sequence, resulted in a significantly higher β-galactosidase enzyme activity (510±50 Miller Units), when compared to the empty vector control (46±5 Miller Units; P<0.01; **Figure 3C**). However, significantly lower enzyme activity was measured with this promoter, when compared to the P_synth(BRE)_ (1350±140 Miller Units; P<0.05) and the P_*hmtB*_ (5000±100 Miller Units; P<0.01; **Figure 3C**) promoters, which both contained a BRE sequence (**Figure 3A**). The P_*mrt*(*M.t*.)_ promoter only resulted in a β-galactosidase activity (65±5 Miller Units) that was comparable and not significantly different to the empty vector negative control in the enzyme assay (45±5 Miller Units; **Figure 3C**). Therefore, this promoter has to be considered inactive under the tested conditions (**Figure 3C**). We did not include the commonly used P_*mcrB*(*M.v*.)_ promoter for methanogen genetic systems in this comparison, because we already had found that this promoter is not functional in driving the neomycin-selectable marker (***SI Appendix*, Results S3**).

### The metabolism of *M. thermautotrophicus* ΔH can be amended to enable growth on formate as an alternative substrate

Formate as the sole carbon and energy substrate can be utilized by several methanogens, such as *Methanococcus* spp. (41), *Methanobacterium* spp., and also *Methanothermobacter* spp. (42). For example, the strain *M. thermautotrophicus* Z-245 can grow with only formate, instead of hydrogen and carbon dioxide, while *M. thermautotrophicus* ΔH is not able to grow with only formate (**Figure 4; *SI Appendix*, Figure S8**) (42, 43). It was hypothesized by Nölling and Reeve (1997) that the genetic reason for this is the missing formate dehydrogenase (*fdh*)-operon in the genome of *M. thermautotrophicus* ΔH when compared to the same genomic region in *M. thermautotrophicus* Z-245. We argued that we can test this hypothesis *in vivo*, by providing the *fdh*-operon as a genetic cargo in the application module of our shuttle vector. Thus, we constructed the shuttle vector pMVS1111A:P_*hmtB*_-*fdh*_Z-245_ that contains the entire *fdh*-operon from *M. thermautotrophicus* Z-245, including the genes (in this order) *fdhC*, *fdhA*, *fdhB*, and additionally an open-reading frame with unknown function (*orf3*), as indicated in Nölling and Reeve (1997), under the control of the constitutive P_*hmtB*_ promoter (**Figure 3**). In control experiments with *M. thermautotrophicus* Z-245, we confirmed growth of this microbe with either formate or hydrogen and carbon dioxide as substrates (***SI Appendix*, Figure S8**). Growth on formate was possible with *M. thermautotrophicus* ΔH cells that carry pMVS1111A:P_hmtB_-*fdh*_Z-245_, but not with cells that carry the empty vector control pMVS-V1 (**Figure 4**). The formate dehydrogenase-producing strain had a prolonged lag-phase with formate, but reached a comparable final OD_600_, when compared to growth on hydrogen and carbon dioxide. Hence, the hypothesis by Nölling and Reeve (1997) was proven to be correct.

**Figure 4.**
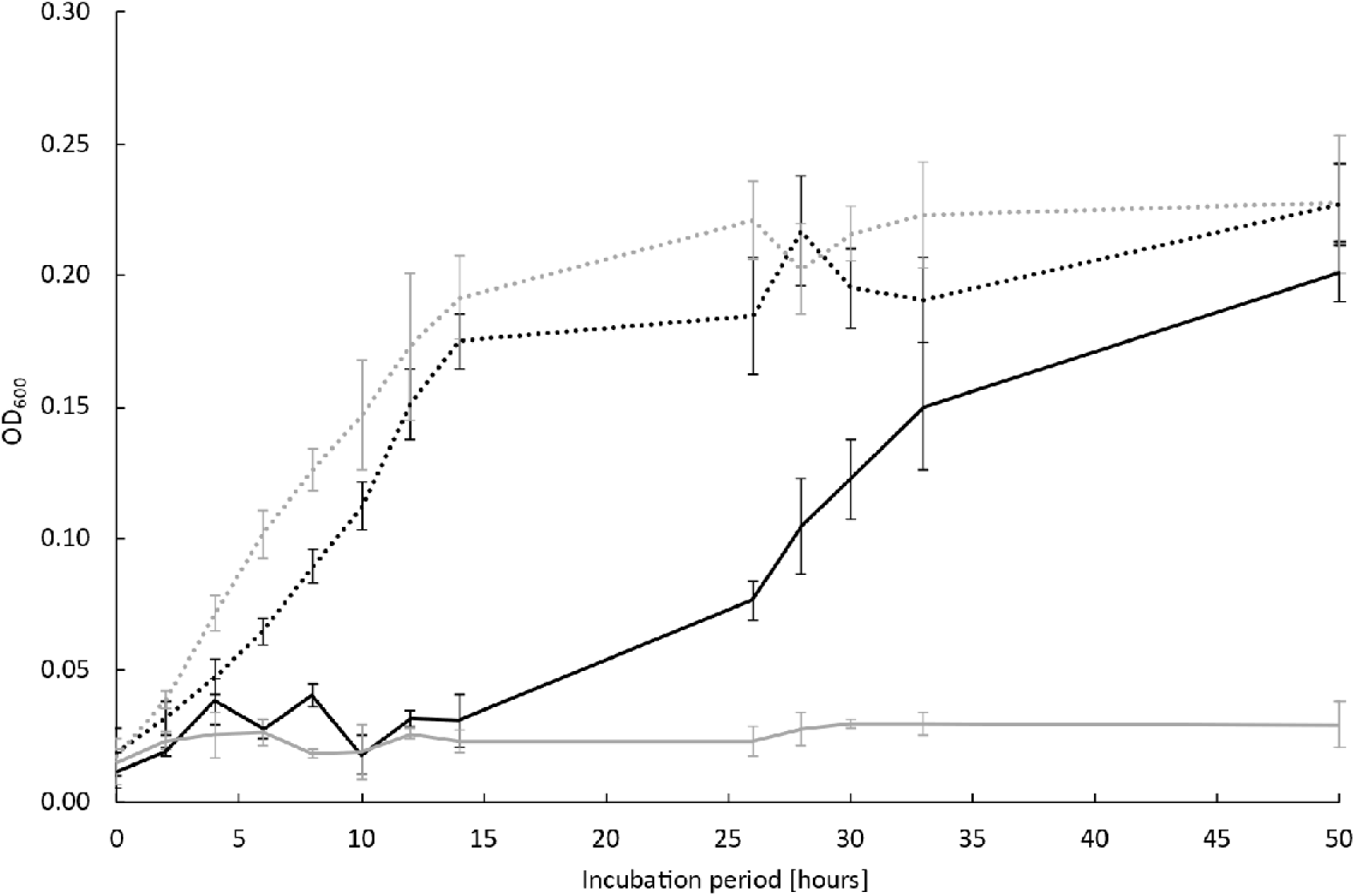
Analysis of genetically modified *M. thermautotrophicus* ΔH strains for growth on formate. Growth of pMVS-V1- (**grey**) and pMVS1111A:P_*hmtB*_-*fdh*_Z-245_- (**black**) carrying *M. thermautotrophicus* ΔH with either hydrogen and carbon dioxide (**dotted line**) or with formate (**solid line**) as the carbon- and energy source. Average (*N*=3) with error bars indicating standard deviation.

## Discussion

Here, we report a robust method for genetic manipulation of *M. thermautotrophicus* ΔH (**Figure 2**), including a modular vector system (pMVS). We demonstrated the function of a thermostable β-galactosidase-encoding gene (*bgaB*) from *G. stearothermophilus* as a reporter to study gene expression, and we implemented a formate dehydrogenase-encoding operon from *M. thermautotrophicus* Z-245 to amend the metabolism of *M. thermautotrophicus* ΔH for growth on formate instead of hydrogen and carbon dioxide.

To achieve this, in a first step, we investigated existing protocols to obtain clonal populations as individual colonies on solidified media plates by using different plating techniques. While we achieved plating efficiencies of up to 100%, the plating conditions, such as humidity of the gas phase, growth phase of the microbial culture, and plating technique, considerably influenced the outcome over a range of three orders of magnitude (***SI Appendix*, Discussion S1**). Thus, for specific purposes the plating technique has to be carefully considered to avoid misleading results, such as false interpretations of DNA-transfer events. At a too high plating efficiency, spontaneous-resistant cells might overgrow genetically modified *M. thermautotrophicus* ΔH, because timing is of utmost importance for successful experiments as further discussed below.

With optimized plating conditions, we were able to confirm growth inhibition of *M. thermautotrophicus* ΔH with antibiotics on solidified media plates. For simvastatin, the inhibiting concentration was 50 μM (21.5 μg/mL), which is higher compared to inhibiting concentrations that were reported to be required for *P. furiosus* or *M. jannaschii* (10-20 μM or 4-8 μg/mL) (26, 34). For neomycin, the inhibiting concentration was 250 μg/mL, which is lower compared to *Methanococcus* spp. for which 500 μg/mL are commonly applied for transformation experiments (16). In general, it was noticeable that spontaneous-resistant *M. thermautotrophicus* ΔH cells appeared readily with both simvastatin and neomycin. Once those spontaneous-resistant cells appeared, they were not inhibited or delayed in growth in subsequent transfers in liquid media as well as on solidified media plates. This has been already reported with neomycin for *M. thermautotrophicus* ΔH (14), *M. maripaludis*, and *Methanococcus vanielli* (22). However, in comparison to *M. maripaludis* for which spontaneous neomycin-resistant colonies appeared to be smaller than genetically modified colonies (which were found to carry a neomycin-selectable marker) (44), we cannot see this difference in colony size, potentially due to apparently faster growth rates of *M. thermautotrophicus* ΔH on solidified media plates. Therefore, we implemented a selective-enrichment step in liquid media, which provides enough time for growth of the genetically modified *M. thermautotrophicus* ΔH, while at the same time excludes the onset of growth of the spontaneous-resistant cells by substrate limitation in the gas phase, because those cells appear only after a longer incubation period. After the selective-enrichment step, the genetically modified *M. thermautotrophicus* ΔH cells outcompeted the spontaneous-resistant cells sufficiently to obtain and select genetically modified individual colonies on selective solidified media plates.

With a slightly modified protocol without the selective-enrichment step and with a prolonged non-selective recovery, we observed conjugation frequencies ranging from 4·10^-9^ to 6·10^-6^, which was only approximately three to ten individual colonies in absolute numbers per solidified media plate in various experiments under the utilized experimental conditions (***SI Appendix*, Results S6**). In our standard protocol, which includes the selective-enrichment step, we found many more individual colonies, which makes the transfer of a shuttle vector efficient and reliable. However, these individual colonies do not represent independent genetically modified *M. thermautotrophicus* ΔH cells, but represent a population, which is derived from only a few transconjugant cells. To further enhance the conjugation frequencies, a reduction of the size of the pME2001 replicon can be considered, as it was shown for the replicon pURB500 in *M. maripaludis* (25). Furthermore, a concentration of *E. coli* and *M. thermautotrophicus* ΔH cells for spot-mating on filter paper might lead to higher conjugation frequencies (45). Also, the ratio of *E. coli* to *M. thermautotrophicus* ΔH cells can be further investigated, as well as other parameters such as the growth conditions during the mating process or different conjugative *E. coli* strains.

With the reporter gene *bgaB*, we tested a set of four distinct promoters (P_*mrt*(*M.t*.)_, P_synth_, P_synth(BRE)_, and P_*hmtB*_) (**Figure 3**). The promoter region upstream of the *mtr* gene of *M. thermautotrophicus* ΔH (P_*mrt*(*M.t*.)_) did not lead to production of a functional BgaB. This is consistent with transcription assays with the *mrt* gene from Darcy *et al*. (1999), where no *in-vitro*-transcription activity with P_*mrt*(*M.t*.)_ could be observed. There is evidence found in the peer-reviewed literature, that this might be because of regulatory effects based on the availability of hydrogen, which might in turn influence the gene expression from the P_*mrt*(*M.t*.)_ promoter (47). In contrast, with the promoters P_synth_, P_synth(BRE)_, and P_*hmtB*_, which are all based on the promoter region upstream of the *hmtB* gene of *M. thermautotrophicus* ΔH, we found significant differences in the β-galactosidase enzyme activity when compared to pMVS-V1 without the *bgaB* gene (**Figure 3C**). The BRE sequence upstream of the P_synth_ sequence, which we had implemented in P_synth(BRE)_, influenced the promoter strength significantly. Similar observations were made with archaeal promoters in *Sulfolobus solfataricus* and *Haloferax volcanii*, respectively, where promoter strength was influenced by the presence or absence of a BRE sequence (48, 49). Nevertheless, the enzyme activity when using the native P_*hmtB*_ was enhanced significantly when compared to both P_synth_ and P_synth(BRE)_ (**Figure 3C**). This is potentially due to modifications in the transcription-initiation region in the promoter sequence, which was also shown to influence the strength of gene expression in *Sulfolobus acidocaldarius* and *S. solfataricus* (48, 50). Further *in-vivo* promoter studies will elucidate more details on the promoter structures of *M. thermautotrophicus* ΔH. For this, the thermostable β-galactosidase as a reporter provides an adequate basis.

In addition to demonstrating heterologous production of functional β-galactosidase enzymes in *M. thermautotrophicus* ΔH, we demonstrated the production of an active formate dehydrogenase enzyme from the *fdh* operon from *M. thermautotrophicus* Z-245, which amended the metabolism of *M. thermautotrophicus* ΔH for the ability to utilize formate as an alternative growth substrate (**Figure 4**). These results are a cornerstone for heterologous and homologous (over)-expression of genes in *M. thermautotrophicus* ΔH to study many hypotheses from four decades of research on a genetic level.

We believe that the most important parameters for the successful implementation of genetic tools for *M. thermautotrophicus* ΔH, when compared to previous attempts during the last four decades, were: **1)** the adaptations to the conjugation protocol, specifically the applied temperature, media, and headspace gas conditions during the spot-mating procedure, in combination with the selective-enrichment step with limited gas supply that facilitated the selection for genetically modified cells over spontaneous-resistant cells; **2)** the utilization of a constitutive native promoter sequence, which was demonstrated to be functional *in-vitro* and in other thermophilic archaea, because the classical P_*mcrB*(*M.v*.)_ promoter turned out to be inactive in *M. thermautotrophicus* ΔH; and **3)** the construction of the shuttle vector with restriction/ligation-independent cloning to fuse the pME2001 replicon with the other modules precisely at the IF5 sequence, to not interrupt any open reading frame or potential sequence of the origin of replication.

With these genetic tools, we can now modify and amend the metabolism of *M. thermautotrophicus* ΔH not only on the substrate, but also on the product side. The possibility to change the product spectrum of hydrogenotrophic methanogens has been demonstrated already for *M. maripaludis* with a genetic modification that resulted in geraniol production (51). Our genetic tools for heterologous gene expression enable us now to broaden the product spectrum of *M. thermautotrophicus* ΔH, and to utilize this industrially relevant and robust microbe for power-to-chemicals or power-to-x applications.

## Materials and Methods

### Microbial strains, media, and cultivation conditions

*M. thermautotrophicus* ΔH (DSM 1053), *M. thermautotrophicus* Z-245 (DSM 3720), and *M. marburgensis* (DSM 2133) were obtained from the DSMZ (Braunschweig, Germany), and were cultivated in mineral medium with modifications to the media composition when required (***SI Appendix, Materials and Methods***). *E. coli* strains for general cloning, conjugation experiments, and heterologous PeiP production, as well as the respective growth conditions are described in ***SI Appendix, Materials and Methods***.

### Molecular cloning and plasmid construction

All primers, gBlock^™^ DNA fragments (IDT, Coralville, IA, USA), and plasmids used in this study are summarized in **Tables S2-S4**. Molecular cloning methods and the detailed cloning strategies to generate the various plasmids and shuttle vectors used in this study are provided in ***SI Appendix, Materials and Methods***.

### Plasmid DNA extraction from *Methanothermobacter* spp

Plasmid DNA from wild-type *M. marburgensis* cells or shuttle vector-carrying *M. thermautotrophicus* ΔH was extracted with the QIAprep Spin Miniprep kit (Qiagen, Hilden, Germany) according to the manufacturer’s guidelines in combination with an enzymatic lysis step with the cell wall-degrading pseudomurein endoisopeptidase PeiP (***SI Appendix, Materials and Methods***).

### Genetic manipulations of *M. thermautotrophicus* ΔH

DNA transfer *via* interdomain conjugation was achieved between *E. coli* S17-1 and *M. thermautotrophicus* ΔH (***SI Appendix, Materials and Methods***). The genetically modified *M. thermautotrophicus* ΔH strains were analyzed *via* sub-cultivation in selective conditions, site-specific PCR amplifications, and plasmid extraction with subsequent retransformation of *E. coli (**SI Appendix, Materials and Methods***). The segregational stability of the shuttle vector pMVS-V1 was analyzed in *M. thermautotrophicus* ΔH *via* sub-cultivation under non-selective conditions in liquid culture and on solidified media plates (spread-plating), and PCR analysis of individual colonies (***SI Appendix, Materials and Methods***). Conjugation frequencies (*i.e*., transconjugants per initial recipient cells) were determined by cell counting of recipient cell cultures in a Petroff-counting chamber and by counting of individual colonies after conjugation experiments with a modified conjugation procedure (***SI Appendix, Materials and Methods***).

### β-Galactosidase enzyme activity assays

For qualitative β-galactosidase enzyme activity assays with genetically modified *M. thermautotrophicus* ΔH strains in liquid cultures, we used S-Gal as the chromogenic substrate (**Figure 3B**). For quantitative β-galactosidase enzyme activity assays, we used ONPG as the chromogenic substrate, following a modified protocol according to Jensen *et al*. (2017). Enzyme activity was defined in Miller Units as change in absorbance at 420 nm per assay time in hours, optical density at 600 nm, and volume of *M. thermautotrophicus* ΔH cell culture (ΔA_420_·(h·OD_600_·L)^-1^) (***SI Appendix, Materials and Methods***).

## Author Contributions

B.M. and L.T.A. initiated the work. C.F. and B.M. designed the experiments. C.F., S.B., A.M.E., and L.M. performed laboratory experiments and analyzed the data. L.T.A. and B.M. supervised the project. C.F. and B.M. wrote the manuscript, while all edited the paper and approved the final version.

## Acknowledgments

The authors are grateful to John Reeve, William W. Metcalf, Rudolf K. Thauer, and Michael Rother for helpful discussions. The authors acknowledge Caroline Schlaiß, Sylvia Lemke, and Gabriela Contreras-Arriagada for performing supportive experiments, Luis Antoniotti from the Max Planck Institute for Developmental Biology workshop for his technical input during the design of the stainless-steel jars, and the Archaea center of the University of Regensburg for kindly providing the plasmids pYS3 (Winfried Hausner) and pME2508. The work was funded by the Alexander von Humboldt Foundation in the framework of the Alexander von Humboldt Professorship (L.T.A.) and the U.S. Office of Naval Research Global (ONRG, N62909-19-1-2076; L.T.A., B.M.). Additional funding sources were the German Federal Ministry of Education and Research (MethanoPEP, 031B0851C; B.M.) and the Deutsche Forschungsgemeinschaft (DFG, German Research Foundation) under Germany’s Excellence Strategy – EXC 2124 – 390838134 (L.T.A., B.M.).

