## Supplementary material for "A shuttle-vector system allows heterologous gene expression in the thermophilic methanogen *Methanothermobacter thermautotrophicus* ΔH": SI Appendix

**This PDF file includes:**

Supplementary Results S1 to S9
Supplementary Discussion S1
Supplementary Figures S1 to S8
Supplementary Tables S1 to S4

### SI Appendix

### SI Results

#### Results S1 – Plating efficiencies

Growth on solidified media plates was investigated with spot-, spread-, and pour-plating (***Materials and Methods***). With spot-plating, colonies are barely distinguishable, but concentrated and dense growth can be obtained (**Figure S1A**). With spread- and pour-plating, we obtained individual colonies (**Figure S1B, C**), and therefore we further investigated factors that influence plating efficiencies with these techniques. While the experimental variance was high in individual experiments and strongly depended on many different factors, as further discussed below, some factors had a distinct impact on the plating efficiency. After we had optimized the plating conditions, we performed a set of experiments to compare the plating efficiency with as little experimental variance as possible (**Figure S1D**). The addition of 0.1 volume% hydrogen sulfide in the headspace gas mixture, as additional reducing agent and sulfur source, resulted in an increase of the number of individual colonies by one order of magnitude when compared to the same plating procedure without hydrogen sulfide gas (**Figure S1D**). With hydrogen sulfide gas and with spread-plating of cells in the stationary growth phase, the plating-efficiency was found to be 1.2±0.5% (*N*=6; **Figure S1D**), while with spread-plating of cells in the exponential growth phase, a plating efficiency of 5±2% (*N*=4) was reached (**Figure S1D**). When we used pour-plating with the same number of cells in the stationary growth phase as with spread-plating, and with hydrogen sulfide gas, the number of individual colonies increased drastically by two orders of magnitude with a plating efficiency of up to 50% (**Figure S3B**) or higher (135±10%, *N*=3; **Figure S1D**), whereas the colony size decreased when compared to colonies from spread-plating (**Figure S1B, C**). However, larger colonies on top of the pour-plated solidified media plates were also observed (**Figure S1B**). Plating efficiencies of more than 100% and high standard deviations in independent experiments appeared from incomplete counting, because of the formation of filaments or clumps of cells, and from a degree of sensitivity of the plating procedure to variations in the experimental handling, such as differences in the exact growth phase of the plated culture or mineral media batches (**Discussion S1**). In many experiments, we realized that the plating efficiency decreased considerably, when water accumulated inside the plate, and formed a layer on the side of the plate, which led to a seal between the bottom of the petri dish-plate and the lid and prevented sufficient gas exchange. As a protective measure, we implemented the addition of paper clips on the edges of the plate to lift up the lid slightly, which efficiently prevented water from sealing the plates (**Figure S1E; Discussion S1**).

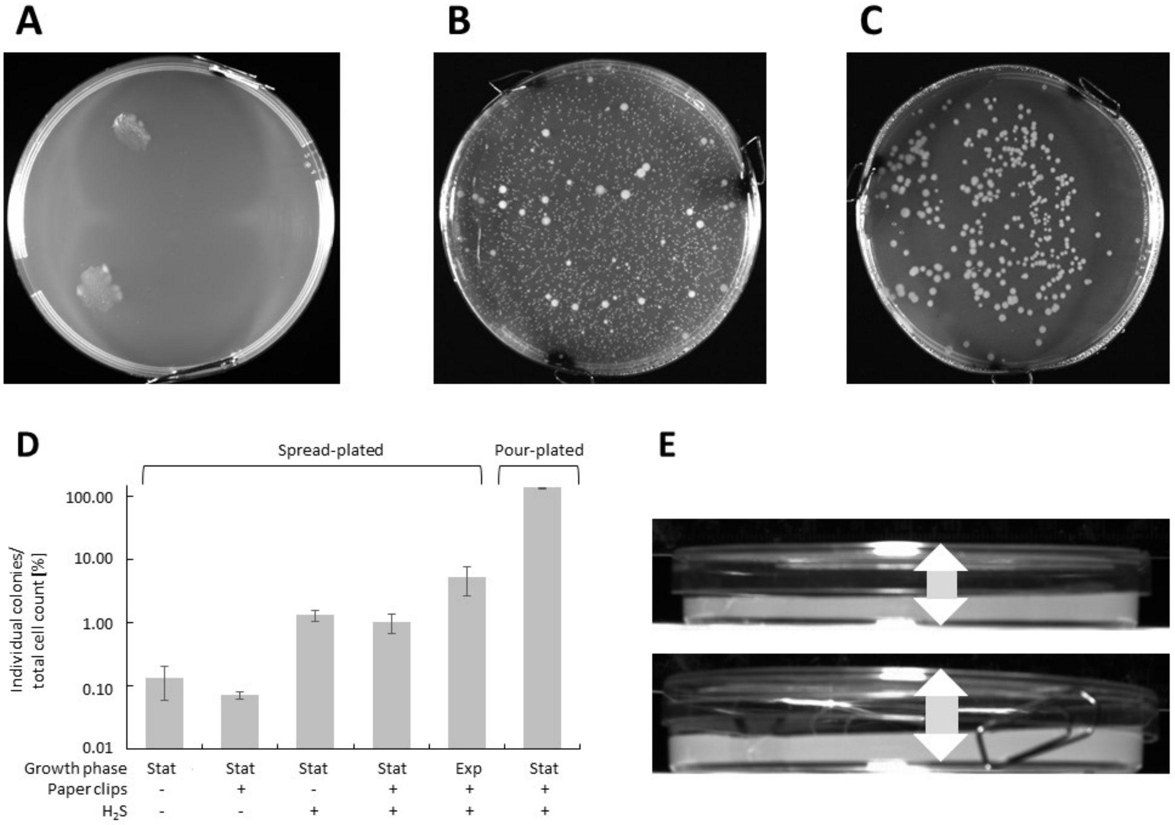


**Figure S1. A)** Spot-plating of *M. thermautotrophicus* ΔH cells on solidified media plates. **B)** Pour-plating of $1{\cdot}10^4$ *M. thermautotrophicus* ΔH cells on solidified media plates. **C)** Spread-plating of $1{\cdot}10^4$ *M. thermautotrophicus* ΔH cells on solidified media plates. **D)** Comparison of the influence from various factors on the plating efficiency with spread-plating of $1{\cdot}10^4$ *M. thermautotrophicus* ΔH cells and pour-plating of $1{\cdot}10^3$ *M. thermautotrophicus* ΔH cells, including the influence of paper clips, 0.1 volume% hydrogen sulfide (**$H_2S$**) in the head-space gas mixture, exponential growth phase (**Exp**), and stationary growth phase (**Stat**). The bars give the average percentage (log-scale) of individual colonies per cell-count of initial microbial cells used for plating from technical replicates (*N*=3; *N*=4 for spread-plating with paper clips and $H_2S$ in exponential growth phase) with error bars indicating standard deviation. **E)** Influence of paper clips on the lid elevation. The arrow demonstrates the increase of lid/plate space compared for a plate without paper clips (upper picture) and with paper clips (lower picture).

### Results S2 – Growth-inhibiting effects of antibiotics

We investigated the growth-inhibiting effects of common antibiotics (simvastatin/mevilonin, neomycin, and puromycin) on *M. thermautotrophicus* ΔH in liquid cultures, which are known to inhibit the growth of methanogens (1-3). We found that non-selective conditions result in a densely grown culture within 24 h. For simvastatin, we tested concentrations ranging from 0-21.5 µg/mL (0-50 µM) on ~$5{\cdot}10^5$

cells/mL, and found that 13 µg/mL (30 µM) inhibited growth at least for 48 h. We also tested the inhibition by the simvastatin-analog mevilonin, but did not find a growth-inhibiting effect on *M. thermautotrophicus* ΔH up to a concentration of 21.5 µg/mL (50 µM). For neomycin, we tested concentrations ranging from 0-250 µg/mL on ~$5 \cdot 10^5$ cells/mL initial cell density. We found that 50 µg/mL inhibited growth for less than 24 h, 100 µg/mL inhibited growth for at least 48 h, and 250 µg/ml neomycin for at least 60 h of incubation, while we did not analyze the growth-inhibiting effect beyond an incubation period of 60 h (**Figure S2**). We further investigated puromycin in liquid mineral medium, because this antibiotic is commonly used in genetic systems for mesophilic methanogens (1, 4), and we found good inhibition from 50 µg/mL on ~$5 \cdot 10^5$ cells/mL initial cell density for at least 72 h. However, the available selectable marker has not yet been adapted to thermophilic conditions to confer resistance against puromycin at elevated temperature. Thus, we did not proceed with this antibiotic.

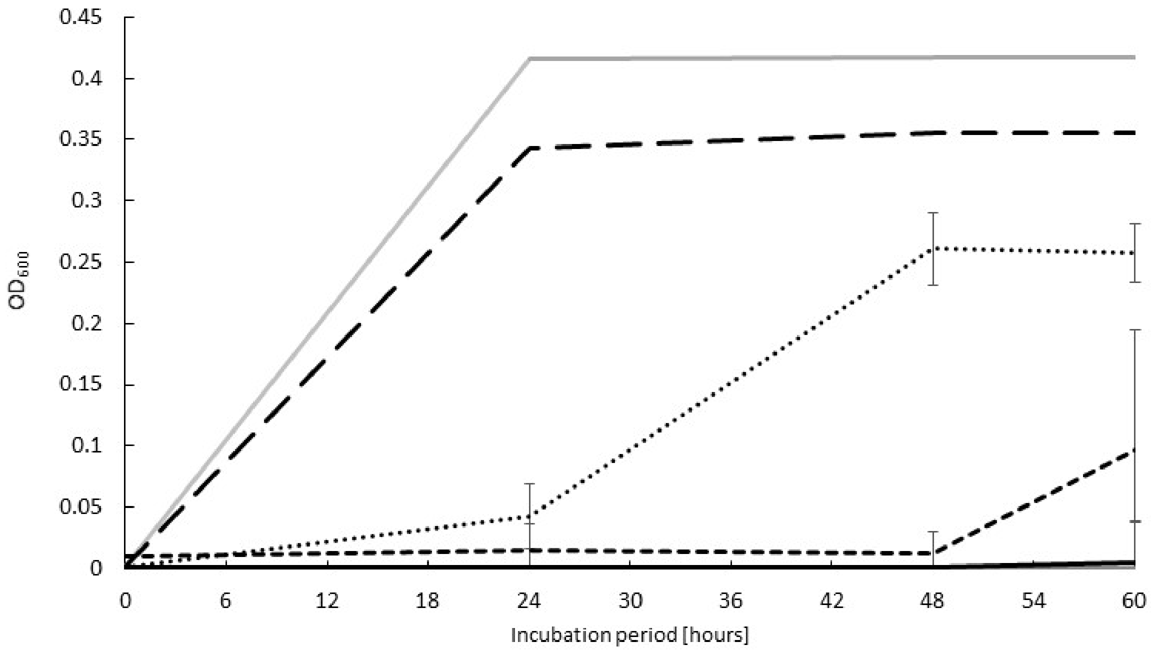


**Figure S2.** Measurement of $OD_{600}$ after different incubation periods at 60°C (0 h, 24 h, 48 h, and 60 h) with initially $5 \cdot 10^5$ cells/mL of wild-type *M. thermautotrophicus* ΔH in liquid mineral medium containing neomycin concentrations of 0 µg/mL (**long-dashed line**, *N*=1), 50 µg/mL (**dotted line**), 100 µg/mL (**short-dashed line**), and 250 µg/mL (**black solid line**). For comparison to wild-type *M. thermautotrophicus* ΔH, initially $5 \cdot 10^5$ cells/mL pMVS-V1-carrying *M. thermautotrophicus* ΔH was incubated in selective liquid mineral medium with 250 µg/mL neomycin (**grey solid line**, *N*=1) and analyzed after the same incubation periods. Average (*N*=3) with error bars indicating standard deviation.

With having the plating efficiency defined (**Results S1**), we further investigated antibiotics for the inhibitory effects on solidified media plates. For simvastatin, we tested concentrations ranging from 0-21.5 µg/mL (0-50 µM) on the inhibition of $1 \cdot 10^8$ cells with pour-plating (**Figure S3A**). While up to 8.7 µg/mL (20 µM) simvastatin resulted in a microbial lawn of *M. thermautotrophicus* ΔH cells, inhibition was recognizable from 13 µg/mL (30 µM) upwards. At 21.5 µg/mL (50 µM) we observed only 27±10 (*N*=3) individual colonies after an incubation period of 48 h (**Figure S3A**), which corresponds to a growth inhibitory efficiency of close to 100%. For neomycin, we tested concentrations ranging from 0-250 µg/mL on the inhibition of $1 \cdot 10^3$ cells with pour-plating (**Figure S3B**). At 50 µg/mL no significant reduction of the number of colonies was observed when compared to non-selective conditions for which we achieved a plating efficiency of ~50% (**Figure S3B**). With 100 µg/mL only 30±5 (*N*=3) individual colonies, and with 250 µg/mL only 3±1 (*N*=3) individual colonies were observed after an incubation period of 48 h, respectively (**Figure S3B**). Therefore, we conclude that 250 µg/mL of neomycin inhibits growth of wild-type *M. thermautotrophicus* ΔH on solidified media plates with an efficiency of >99.9% for at least 48 h (**Figure S3B**). We further tested a genetically modified *M. thermautotrophicus* ΔH strain, which carries pMVS-V1 (**Figure 1**), under selective conditions in liquid mineral medium and on solidified media plates. We found that pMVS-V1 relieves the growth-inhibiting effect of 250 µg/mL neomycin, and resulted in wild-type-like growth (**Figure S2 and S3B**).

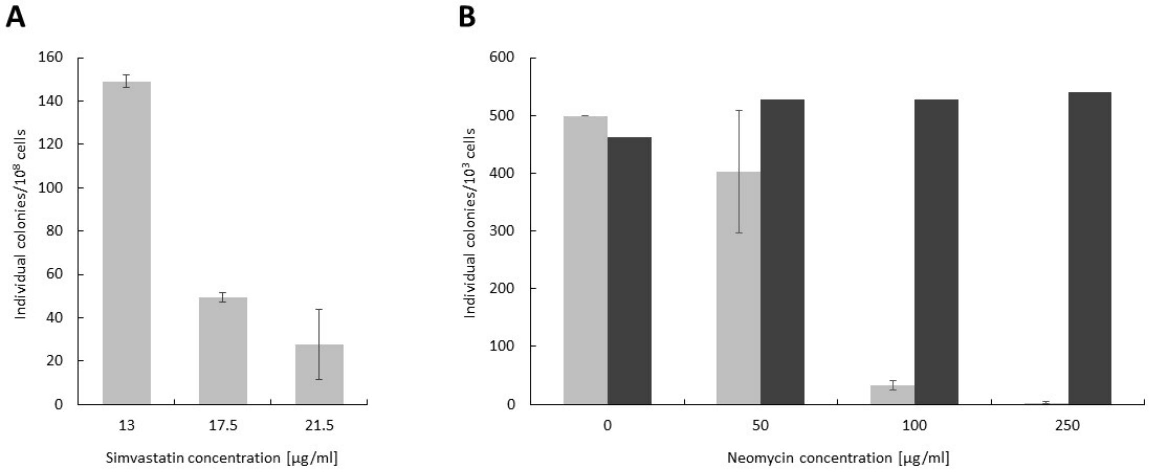


**Figure S3. A)** Individual colonies from $1 \cdot 10^8$ wild-type *M. thermautotrophicus* ΔH cells (**grey bars**), which were pour-plated on solidified media plates containing simvastatin concentrations ranging from 13 µg/mL to 21.5 µg/mL. **B)** Individual colonies from $1 \cdot 10^3$ wild-type *M. thermautotrophicus* ΔH cells (**grey bars**) and pMVS-V1-carrying *M. thermautotrophicus* ΔH (**black bars**, *N*=1), which were pour-plated on solidified media plates containing neomycin concentrations ranging from 0 µg/ml to 250 µg/mL. Average (*N*=3) with error bars indicating standard deviation for wild-type *M. thermautotrophicus* ΔH.

**Results S3 – pMVS design**

The five modules of the pMVS design are separated by rare eight base-pair recognition sequences as described in the main manuscript (*Pme*I, *AsiS*I, *Fse*I, *Asc*I, and *Pac*I). The archetype shuttle vector for this design is pMVS-V1, which does not contain a *Pac*I site (**Figure 1**). We selected the pMTL83151 plasmid from the pMTL80000 system (5) as the source for the backbone for *E. coli* in pMVS-V1, because this plasmid contains the *tra*-region from RK2 (for mobilization of the plasmid) in addition to the ColE1 replicon. Furthermore, this plasmid already brings a chloramphenicol-selectable marker ($Cam^r$) for selection in *E. coli*, which is separated from the replicon by a *Pme*I-recognition sequence, due to the modularity of the pMTL80000 system (5). As the replicon for *M. thermautotrophicus* ΔH, we chose the cryptic plasmid pME2001 from *M. marburgensis* (6). This plasmid has been studied to some extent and is the smallest plasmid, which is known in *Methanothermobacter* spp. (7-9). The plasmid contains five open-reading frames (*orf1-5*) with barely annotated functions. Additionally, in a ~1-kilobase section on pME2001, no open-reading frames have been annotated, but instead this section contains five sites of inserted fragments (IFs), which are different in pME2001 when compared to the similar cryptic plasmid pME2200 from *M. thermautotrophicus* ZH3, whereas IF5 only occurs in pME2001 (7). Previous attempts to create *E. coli*-*M. thermautotrophicus* shuttle vectors relied on the availability of restriction enzyme-recognition sequences in pME2001 (8). To not interrupt one of the open-reading frames and simultaneously to not intersect the region of a potential origin of replication for which the exact location is not known in pME2001 and pME2200, we decided to fuse the pME2001 replicon to the other components of the shuttle vector at the position of IF5 *via* Gibson® Assembly (***Materials and Methods***), and to use the entire plasmid pME2001 as the replicon for *M. thermautotrophicus* ΔH.

We chose the thermostable neomycin-selectable marker ($Neo^r$) from pMU131 for positive selection in *M. thermautotrophicus* ΔH (10). As a promoter, we selected a sequence published by Santangelo *et al.* (2008), which we designate $P_{synth}$ (but which is called $P_{hmtB}$ in Santangelo *et al.* (2008), **Figure 3**), because the commonly used promoter $P_{mcrB(M.v.)}$ from *M. voltae* (12) did not result in genetically modified *M. thermautotrophicus* ΔH in combination with $Neo^r$ in our hands. The promoter sequence from Santangelo *et al.* (2008) has similarity to the upstream region of a histone-binding protein (HmtB)-encoding gene from *M. thermautotrophicus* ΔH. However, several modifications had been introduced in $P_{synth}$ compared to the native $P_{hmtB}$ sequence (13). The native $P_{hmtB}$ sequence was demonstrated to initiate *in-vitro* transcription, and thus, the generation of mRNA when using purified native *M. thermautotrophicus* ΔH RNA polymerase (14). Additionally, the $P_{synth}$ promoter was shown to initiate gene expression in *T. kodakarensis* (11). As the terminator sequence for the selectable marker, we implemented the $T_{mcr}$ terminator sequence of the methyl-coenzyme M reductase (*mcr*) operon from *M. voltae*, which is commonly used in constructs for genetic modification of *Methanococcus* spp. and *Methanosarcina* spp. (1). We selected a thermostable β-galactosidase reporter (**Figure 3**) as our first gene of interest for the application module. When we introduced the thermostable β-galactosidase-encoding gene to pMVS-V1, which resulted in the shuttle vector pMVS1111A:$P_{synth}$-*bgaB* with five modules, we included a *Pac*I site in addition, to complete the application module. The directionality of the *bgaB* gene was such that the $T_{mcr}$ terminator sequence from the selectable marker module is also terminating transcription of the *bgaB* gene (**Figure 1B; *Materials and Methods***).

**Results S4 – Analysis of successful DNA transfer *via* site-specific PCR**

We analyzed genetically modified *M. thermautotrophicus* ΔH strains *via* site-specific PCR amplifications with: **1)** a primer combination (**Table S2**), which specifically amplifies a 1-kilobase fragment of the pME2001 replicon; and **2)** primer combinations (**Table S2**), which specifically amplify either a 1.5-kilobase or a 2.8-kilobase fragment of genomic DNA from *M. thermautotrophicus* ΔH to confirm the integrity of the genetically modified strains (**Figure S4; *Materials and Methods***). In preliminary experiments with high densities of *E. coli* S17-1, we found that PCR signals for the presence of our high-copy number shuttle vectors (with regards to copy number in *E. coli*) could be obtained after up to three transfers in liquid media, and also from areas of solidified media plates at which no growth was observed, after up to two transfers. Therefore, to gather reliable results on the stable replication of shuttle vectors in *M. thermautotrophicus* ΔH, and to exclude false positive results from residual *E. coli* DNA, we first transferred cell material from individual colonies of putative *M. thermautotrophicus* ΔH transconjugants into selective liquid mineral medium. This enrichment culture was plated by streaking some culture with an inoculation loop on selective solidified media plates. From these plates, we inoculated another selective liquid mineral medium with cell material from individual colonies, and these liquid enrichment cultures were analyzed by PCR amplification. Successful DNA transfer into *M. thermautotrophicus* ΔH was reproducibly confirmed (**Figure 2C**).

### Results S5 – Retransformation of *E. coli* with plasmid extracts from *M. thermautotrophicus* ΔH

As a second approach to confirm the presence and integrity of the shuttle vector in *M. thermautotrophicus* ΔH, we extracted plasmid DNA from genetically modified *M. thermautotrophicus* ΔH cultures, and used this plasmid DNA for retransformation of *E. coli* NEB stable (***Materials and Methods***). Individual colonies of *E. coli* NEB stable were analyzed with restriction-enzyme digestion and Sanger sequencing. We found that all analyzed colonies contained the entire shuttle vector as deduced from the correct fragment sizes in the restriction-enzyme digestion (**Figure S4A**), and no mutations in the *M. thermautotrophicus* ΔH replicon as well as the neomycin-selectable marker (**Figure S4B**). We did not re-sequence the entire *E. coli* replicon and selectable marker (**Figure S4B**).

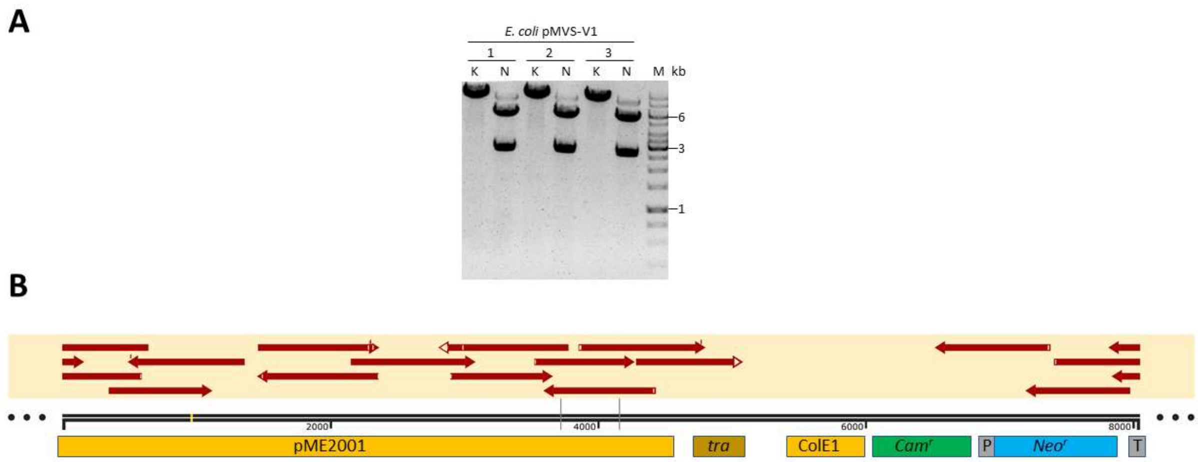


**Figure S4. A)** Restriction-enzyme digestion of plasmid DNA, which was extracted from three independent *E. coli* pMVS-V1 enrichment cultures (1-3). The *E. coli* strains were generated with plasmid DNA, which was extracted from genetically modified *M. thermautotrophicus* ΔH. The unique cutter *Kpn*I (K), resulting in one 8.2-kilobase fragment, and the dual cutter *Nde*I (N), resulting in 2.7-kilobase and 5.5-kilobase fragments were used. **M,** GeneRuler 1 kb DNA Ladder (Thermo Scientific, Waltham MA, USA). **B)** Exemplified sequence alignment of Sanger sequences from retransformed *E. coli* plasmid DNA to the original pMVS-V1 sequence (main text and **Figure 1** for further explanations). Red arrows indicate the sequence and its direction. No deletions, insertions, or nucleotide exchanges were detected.

**Results S6 – Conjugation frequency**

The determination of the conjugation frequency, which we define as transconjugants per initial recipient cells, is accompanied by several unknown parameters, such as the different incubation periods during the spot-mating, and the non-selective-recovery and selective-enrichments steps, before the selective spread-plating step to obtain individual colonies (**Figure 3**). This method reliably resulted in a high number of individual colonies. However, to quantify the conjugation frequency more accurately, we performed a subset of experiments in which we did not include the selective-enrichment step, but instead we prolonged the incubation period for the non-selective-recovery step. Still we had to take different parameters into account, and we corrected the conjugation frequency according to **Equation S1**:

$$f = \frac{N_P \cdot V_R}{E \cdot N_0 \cdot R_W \cdot 2^{(D_S+D_R)}} \quad \textbf{(Equation S1)}$$

where **f** represents the conjugation frequency; $\mathbf{N_P}$ represents the number of individual colonies obtained after the final selective spread-plating step; $\mathbf{V_R}$ represents the dilution factor for the amount of plated cells in relation to overall culture volume; **E** represents the spread-plating efficiency as a fraction; $\mathbf{N_0}$ represents the initial recipient cell number used for spot-mating; $\mathbf{R_W}$ represents the fraction of cells that are recovered from the washing step after the spot-mating; $\mathbf{D_S}$ represents the number of cell divisions during spot-mating; and $\mathbf{D_R}$ represents the number of cell divisions during non-selective recovery.

We calculated a range of parameters in a simple sensitivity analysis assuming the worst and best case scenarios (**Table S1**). The parameters $\mathbf{V_R}$, $\mathbf{N_P}$, and $\mathbf{N_0}$ are either known (dilution factor $\mathbf{V_R}$ is 50 in our experiments, ***Materials and Methods***), or can be experimentally determined by cell counting before the conjugational DNA transfer and colony counting of transconjugants after the selective spread-plating step. The parameter **E** can be experimentally defined with a relatively good certainty, based on our plating efficiencies (**Figure S1**). However, the parameters $\mathbf{R_W}$, $\mathbf{D_S}$, and $\mathbf{D_R}$ need to be estimated in a certain range for the sensitivity analysis, for example, based on assumptions that are made from other experiments. For $\mathbf{D_S}$ this is because in the spot-mating step, *E. coli* and *M. thermautotrophicus* ΔH cells are combined and potential growth (number of cell divisions) of *M. thermautotrophicus* ΔH at 37°C cannot be determined. For our estimations, we define $\mathbf{D_S}$ as either 0 or 1 cell divisions. For $\mathbf{R_W}$, the washing step after the spot-mating step is critical. While with careful suspension of the entire spot a recovery of all cells can be assumed, for our estimations, we define **$R_W$** between 0.5 and 1.0 (for 50-100% recovery of cells). For **$D_R$**, the overall number of cell divisions during the non-selective recovery step is critical. Based on typical growth experiments similar to these conditions (16-20 h without selection), we assume **$D_R$** to be between 5 and 8 cell divisions. These assumptions result in an estimated conjugation frequency between $6 \cdot 10^{-6}$ and $4 \cdot 10^{-9}$ (**Table S1**).

**Table S1.** Example calculations for conjugation frequencies for DNA transfer into *M. thermautotrophicus* ΔH based on Equation S1

| Parameters | $N_P$ | E | $V_R$ | $D_R$ | $N_0$ | $R_W$ | $D_S$ | Conjugation frequency |
|---|---|---|---|---|---|---|---|---|
| Best case scenario | 10 | 0.01 | 50 | 5 | $5 \cdot 10^8$ | 0.5 | 0 | $6 \cdot 10^{-6}$ |
| Worst case scenario | 1 | 0.05 | 50 | 8 | $5 \cdot 10^8$ | 1 | 1 | $4 \cdot 10^{-9}$ |
| Most likely scenario | 8 | 0.02 | 50 | 6 | $5 \cdot 10^8$ | 0.8 | 0 | $8 \cdot 10^{-7}$ |

### Results S7 – Segregational stability of shuttle vector under non-selective conditions

The segregational stability of shuttle vectors in *M. thermautotrophicus* ΔH is of interest for incubation periods under non-selective growth conditions (*e.g.*, in bioreactors), but also for potential plasmid-curing requirements. Therefore, we performed an experiment to assess the segregational stability of pMVS-V1. We inoculated both selective and non-selective liquid mineral medium with ~$5 \cdot 10^5$ cells/mL from a selectively grown pre-culture of pMVS-V1-carrying *M. thermautotrophicus* ΔH. After growth to ~$1 \cdot 10^8$ cells/mL, which corresponds to ~7-8 cell divisions (**Equation S2**), we repeated the transfer of ~$5 \cdot 10^5$ cells/mL from the non-selective liquid mineral medium to non-selective conditions twice. This resulted, in total, in ~21-28 cell divisions under non-selective conditions. We calculated the number of cell divisions according to **Equation S2**:

$$n = \frac{log(N_t) - log(N_0)}{\log(2)} \quad \textbf{(Equation S2)}$$

where **n** is the number of generations (cell divisions), $\mathbf{N_t}$ is the cell concentration at the end of the incubation period, and $\mathbf{N_0}$ is the cell concentration in the beginning of the incubation period.

We spread-plated cells from the first selective transfer to selective solidified media plates, and from all three non-selective transfers to non-selective solidified media plates, respectively. We analyzed 16 individual colonies each from the selective plate (plated from selective liquid transfer) and non-selective plates (plated from all non-selective liquid transfers) *via* site-specific PCR, and found that all analyzed colonies still contained the shuttle vector, independent of the presence of a selection pressure or not (**Figure S5**).

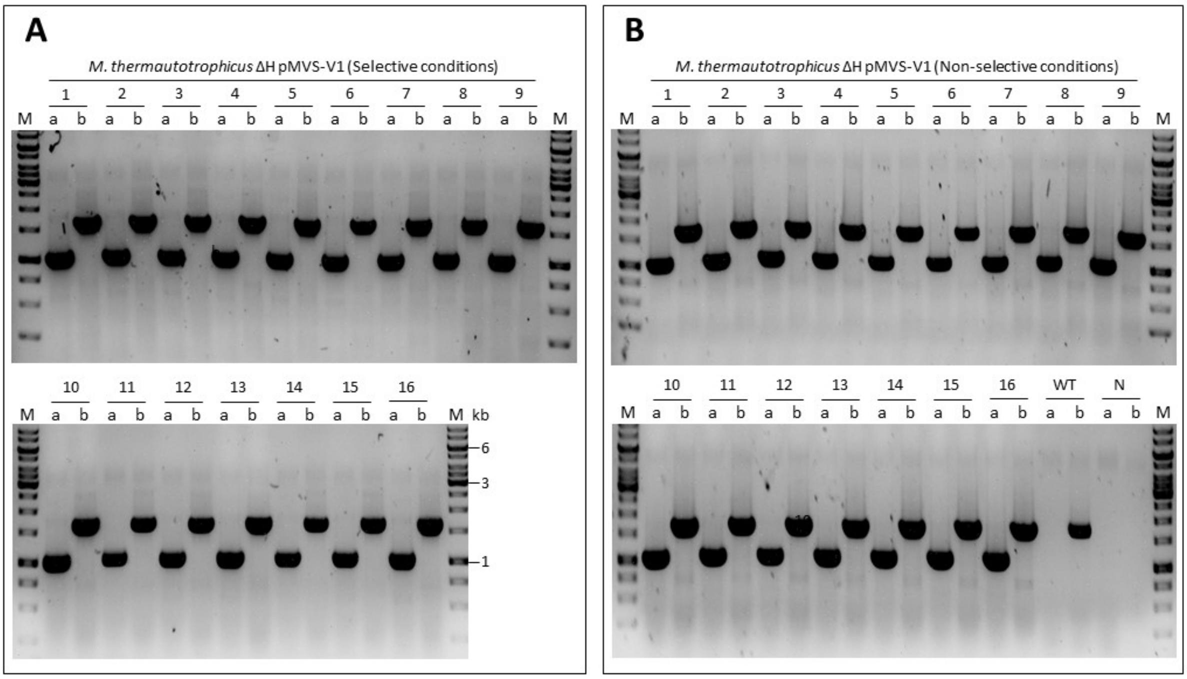


**Figure S5. A)** PCR analysis of individual colonies from selective solidified media plates after one selective transfer of pMVS-V1-carrying *M. thermautotrophicus* ΔH in liquid mineral medium. **B)** PCR analysis of individual colonies from non-selective solidified media plates after **three** non-selective transfers (~21-28 cell divisions) in liquid mineral medium (PCR analysis of the first and second transfer is not shown, but gave the same results). A 1-kilobase fragment results from the specific primer pair for the pME2001 replicon for *M. thermautotrophicus* ΔH **(a)** and a 1.5-kilobase fragment from the specific primer pair for *M. thermautotrophicus* ΔH genomic DNA **(b)**. All 16 colonies each result in positive PCR for both primer combinations. Wild-type *M. thermautotrophicus* ΔH **(WT)** does not result in PCR signal for pMVS-V1, and water as negative control **(N)** does not result in any PCR signal. **M**, GeneRuler 1 kb DNA Ladder (Thermo Scientific, Waltham MA, USA).

### Results S8 – Control experiments for conjugational DNA transfer

To investigate whether conjugational DNA transfer leads to plasmid-carrying *M. thermautotrophicus* ΔH or whether free plasmid DNA can result in genetically modified *M. thermautotrophicus* ΔH, we conducted several control experiments in addition to our standard conjugation protocol (***Materials and Methods***). First, we added 200 ng/mL of purified pMVS-V1 plasmid DNA (extracted from *E. coli* NEB stable) to a freshly inoculated non-selective *M. thermautotrophicus* ΔH culture (~$1 \cdot 10^5$ cells/mL). After growth of this culture to the stationary growth phase, to allow for potential natural competence to occur during all growth phases, we treated the cells similar to our standard conjugation protocol, including the non-selective recovery and selective-enrichment steps, and selective spread-plating, but without the spot-mating/plating step. Second, we substituted *E. coli* S17-1 with the non-conjugative *E. coli* NEB stable, which carried pMVS-V1, to analyze the importance of mobilization for DNA transfer into *M. thermautotrophicus*. Third, we heated *E. coli* S17-1, which carried pMVS-V1, to 60°C for 20 min *prior* to spot-mating, to check whether *E. coli* needs to be viable to mediate the DNA transfer into *M. thermautotrophicus* ΔH. Finally, and fourth, we added 250 U/mL DNase I to the *E. coli* S17-1 pre-culture during the last 30 minutes of incubation at 37°C, to reduce the amount of initial free DNA during the spot-mating process.

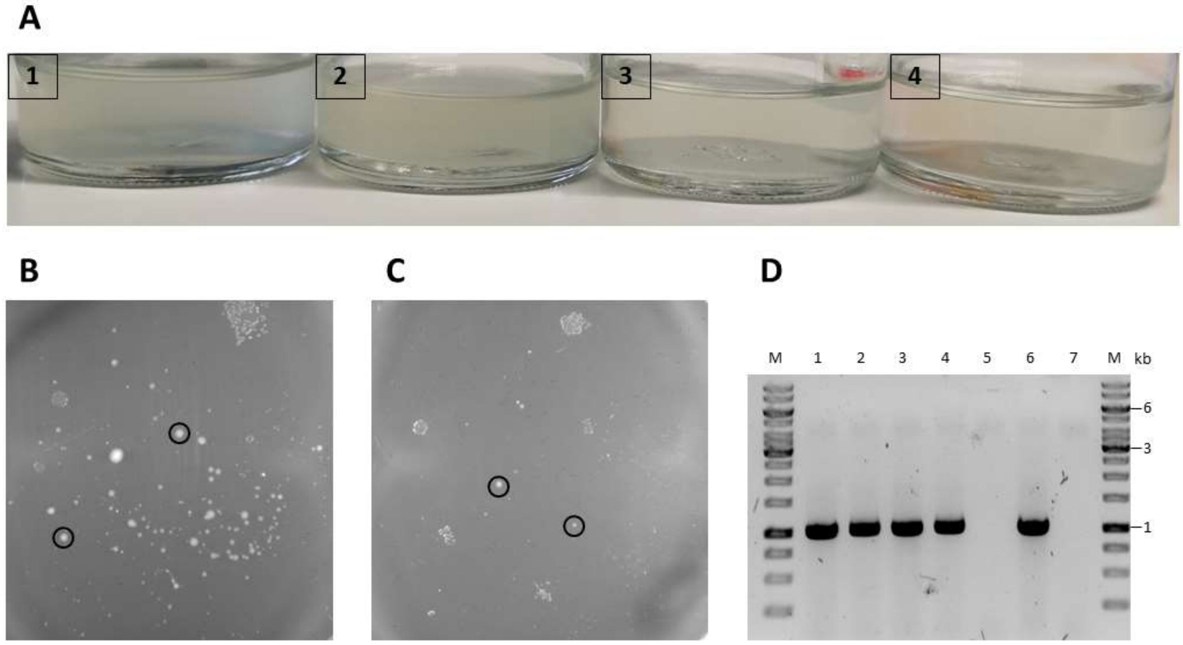


**Figure S6. A)** Enrichment cultures of genetically modified *M. thermautotrophicus* ΔH in 250 µg/ml neomycin-containing selective liquid mineral medium from conjugation/DNA-transfer experiments with the standard protocol **(1)**, the DNase-I treated *E. coli* S17-1 **(2)**, the heat-inactivated *E. coli* S17-1 **(3)**, and NEB stable as non-conjugative *E. coli* **(4)**. Slight turbidity in 3 and 4 is caused due to high initial cell count. Enrichment is visible in 1 and 2. **B)** Spread-plated *M. thermautotrophicus* ΔH using culture 1 from A (standard protocol). Black circles represent colonies, which were used for PCR analysis. **C)** Spread-plated *M. thermautotrophicus* ΔH using culture 2 from A (DNase-I treatment). Black circles represent colonies, which were used for PCR analysis. **D)** PCR analysis of the four colonies from B and C **(1-4)**, wild-type *M. thermautotrophicus* ΔH **(5)**, purified shuttle-vector DNA as positive control **(6)**, and water as negative control **(7)**. A primer combination, which amplifies a 1-kilobase fragment from the pME2001 replicon was used for PCR amplification.

**Results S9 – Quantitative β-galactosidase enzyme activity assay**

While we did not intend to perform an exhaustive promoter study here, the four selected promoters represent a subset of promoter sequences with distinctive features. The first promoter, $P_{synth}$, is a synthetic promoter, which was based on the promoter sequence of the HmtB-encoding gene from *M. thermautotrophicus* ΔH ($P_{hmtB}$) (13), but with several sequence modifications between the start-codon and the TATA-box sequence (**Figure 3A**) (11). This $P_{synth}$ promoter was shown to be functional in *T. kodakarensis* (11), and we had successfully used $P_{synth}$ to drive our neomycin-selectable marker (**Figure 1**). However, we realized after this finding that we had misinterpreted the annotation of this promoter in Santangelo *et al.* (2008), and our $P_{synth}$ promoter did not contain the transcription factor B recognition element (BRE) sequence of archaeal promoters (15). Therefore, as a second promoter, we added the BRE sequence to the $P_{synth}$ promoter sequence to generate the $P_{synth(BRE)}$ promoter sequence, but left all modifications from Santangelo *et al.* (2008) intact (**Figure 3A**). In addition, as the third promoter, we used the promoter sequence upstream of the *hmtB*-gene from *M. thermautotrophicus* ΔH to include the native version of the promoter (13) on which $P_{synth}$ and $P_{synth(BRE)}$ were based (**Figure 3A**). Finally, as the fourth promoter, we included the promoter sequence from upstream of one of the two isoforms for the methyl-coenzyme M reductase operons from *M. thermautotrophicus* ΔH, the $P_{mrt(M.t.)}$ promoter (**Figure 3A**). The $P_{mrt(M.t.)}$ promoter does not contain the typical BRE and TATA-box sequences of archaeal promoters, but instead palindromic sequences that consist exclusively of thymine and adenine bases (16) (**Figure 3A**).

For a quantitative β-galactosidase enzyme activity assay, we used the chromogenic substrate ONPG (***Materials and Methods***). To implement the β-galactosidase enzyme activity assay for *M. thermautotrophicus* ΔH, we determined the β-galactosidase activity in different genetically modified *M. thermautotrophicus* ΔH strains after different incubation periods over the course of a growth experiment (**Figure S7**). Samples to assess β-galactosidase activity were taken after four different incubation periods during the growth experiment (subscript number gives time of incubation period in hours): $\mathbf{t_{15}}$, mid-exponential growth phase; $\mathbf{t_{19}}$, late-exponential growth phase; $\mathbf{t_{23}}$, early-stationary growth phase; and $\mathbf{t_{36}}$, late-stationary growth/death phase (**Figure S7**). We determined the β-galactosidase activity after these various incubation periods for four different genetically modified *M. thermautotrophicus* ΔH strains, which carry the empty vector pMVS-V1 as negative control, the pMVS1111A:$P_{synth}$-*bgaB*, or one of two shuttle vectors, pMVS1111A:$P_{\mathit{hmtB}}$-*bgaB* and pMVS1111A:$P_{\mathit{mrt}(M.t.)}$-*bgaB*, which have the $P_{synth}$ promoter exchanged for the $P_{\mathit{hmtB}}$ or the $P_{\mathit{mrt}(M.t.)}$ promoter, respectively. The $P_{synth(BRE)}$ promoter has not yet been investigated with this experiment. We found similar growth behaviors, with similar maximum $OD_{600}$ values of ~0.4, for all four strains, and only the empty vector pMVS-V1-carrying strain reached the late-exponential growth phase already after 19 h, instead of after 23 h for the other three strains (**Figure S7A-D**). The β-galactosidase activity, given as Miller Units (***Materials and Methods***), was low, with 15±5 and 18±5 Miller Units for the empty vector control (pMVS-V1) and for the pMVS1111A:$P_{\mathit{mrt}(M.t.)}$-*bgaB*-carrying strain after all incubation periods (**Figure S7E**), respectively. For the pMVS1111A:$P_{synth}$-*bgaB*-carrying strain, the activity increased over the course of the growth experiment to 125±15 Miller Units after 36 h of incubation (**Figure S7E**). The pMVS1111A:$P_{\mathit{hmtB}(M.t.)}$-*bgaB*-carrying strain showed the highest β-galactosidase activity with 270±20 Miller Units, while already after 19 h of incubation, no further increase in the enzyme activity was observed (**Figure S7E**).

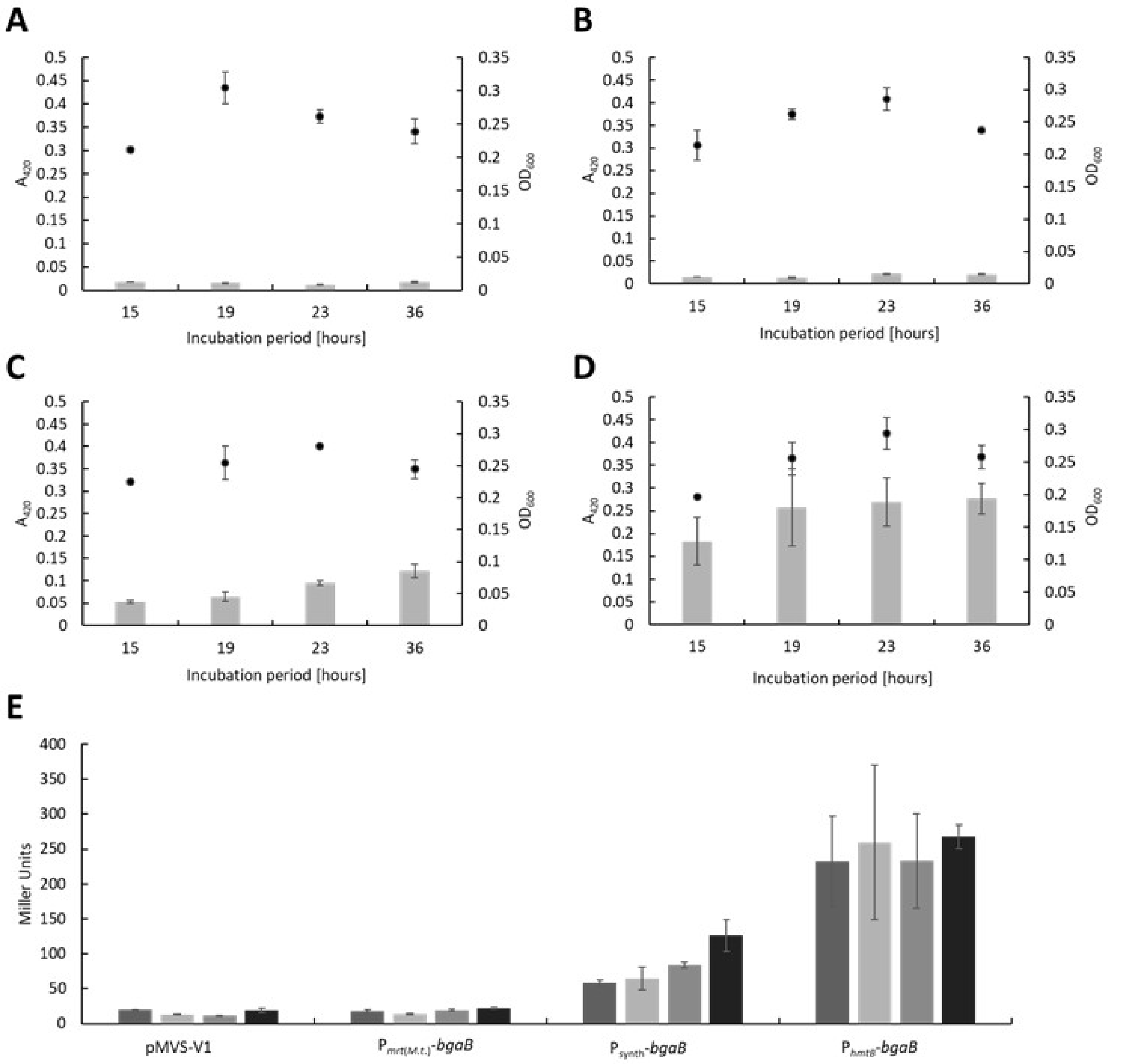


**Figure S7. A-D)** The measurements of absorbance at 420 nm **($A_{420}$)** of the enzyme activity assay **(bars)** and optical density at 600 nm **($OD_{600}$)** of the corresponding genetically engineered *M. thermautotrophicus* ΔH strain **(dots)** after different incubation periods (*M. thermautotrophicus* ΔH with **A**, pMVS-V1; **B,** pMVS1111A:$P_{mrt(M.t.)}$-*bgaB*; **C,** pMVS1111A:$P_{synth}$-*bgaB*; **D,** pMVS1111A:$P_{hmtB}$-*bgaB*). **E)** The resulting Miller Units from A-D for the four different *M. thermautotrophicus* ΔH strains is given for the incubation periods 15 h, 19 h, 24 h, and 36 h for each strain from left to right. Average (pMVS-V1 (*N*=2), pMVS1111A:$P_{mrt(M.t.)}$-*bgaB* (*N*=3), pMVS1111A:$P_{synth}$-*bgaB* (*N*=2), and pMVS1111A:$P_{hmtB}$-*bgaB* (*N*=3)) with error bars indicating standard deviation. The Miller Units in this experiment are lower compared to the experiment in Figure 3C (main manuscript) because of differences in the experimental parameters (Materials and Methods).

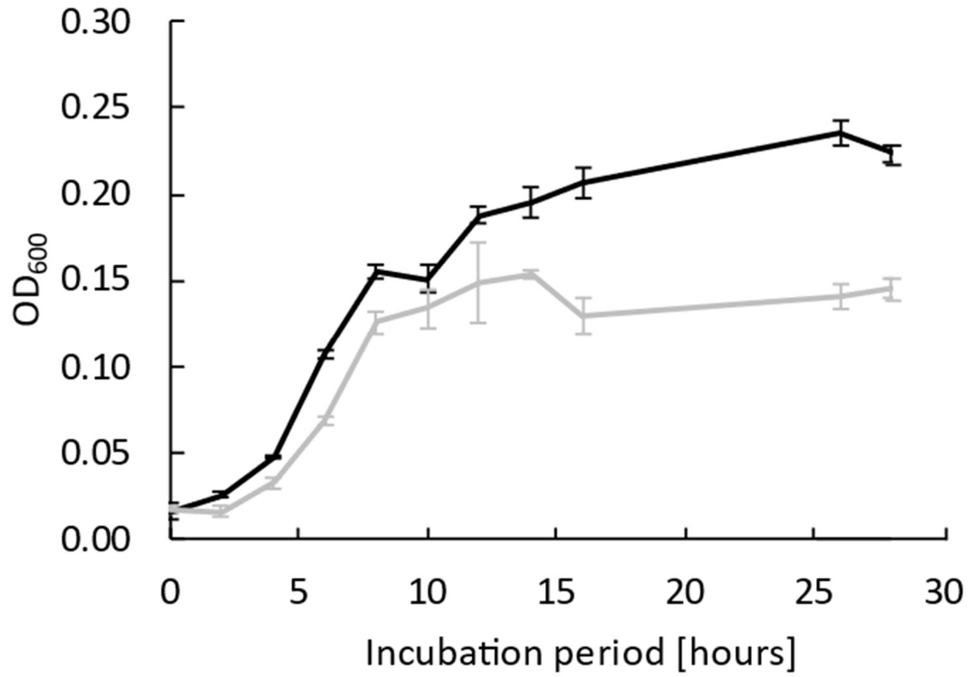


**Figure S8**. Analysis of wild-type *M. thermautotrophicus* Z-245 growth with either hydrogen and carbon dioxide (**black solid line**) or with formate (**grey solid line**) as the carbon- and energy source. Average (*N*=3) with error bars indicating standard deviation.

### SI Discussion

#### Discussion S1 – Plating of *M. thermautotrophicus* ΔH

In several experiments, we observed plating efficiencies of more than 100%. *Methanothermobacter* spp., including *M. thermautotrophicus* ΔH, form filaments of cells and/or show aggregation of cells depending on the growth phase and culture conditions, as has been reported before (17, 18). This complicates cell counting in the Petroff-counting chamber and can result in an underestimation of the initial number of cells for plating, which leads to calculated plating efficiencies of more than 100%. The number of individual colonies on solidified media plates was further dependent on factors, such as the position of the plate within our anaerobic stainless-steel jar. In a typical experiment, up to ten plates were incubated in one jar, and even with plates that were prepared from the same media batch, and with the same liquid *M. thermautotrophicus* ΔH culture, we observed large variations in the number of individual colonies. Furthermore, we found considerably decreased plating efficiencies, when solidified media plates were not sufficiently dry to absorb the *M.*

*thermautotrophicus* ΔH culture completely during the plating procedure. The accumulation of too much water can result from the formation of metabolic water during hydrogenotrophic methanogenesis ($4\,H_2 + CO_2 -> CH_4 + 2\,H_2O$), but also due to incomplete drying of media plates after pouring hot media inside the anaerobic chamber (***Materials and Methods***). Therefore, a drying step was crucial, while this drying step should not exceed two hours in our experimental set-up, to avoid loss of carbon dioxide from the carbonate buffer system in the solidified media plates (because we used a nitrogen atmosphere in our anaerobic chamber; ***Materials and Methods***). While this was sufficient as long as the humidity of the anaerobic chamber was sufficiently low, with accumulation of moisture inside the chamber, a two hour-drying step still resulted in accumulation of water in the plates. Thus, we had implemented paper clips as spacers for the petri dishes (**Figure S1E**), which helped to overcome this issue, especially when the humidity in the anaerobic chamber was high. These paper clips ensured efficient gas-solid mass transfer of hydrogen and carbon dioxide, however, highest plating efficiencies were observed with sufficiently dried solidified media plates. Furthermore, the growth phase of the culture that was used for plating had a considerable effect on colony formation, especially for spread-plating (**Results S1**). A possible reason for this difference could be that cells are more evenly distributed during the pour-plating procedure in comparison to the spread-plating procedure, which might lead to a better accessibility for *M. thermautotrophicus* ΔH to media components, or more stable pH conditions.

### SI Materials and Methods

**Table S2.** List of primers used in this study

| Name | Purpose | Sequence (5’ -> 3’) | Reference |
|---|---|---|---|
| Gib_CF1 | pCF203 | ATAAAATGCTTGGGAGATGACGCCCGCCCCAC | This study |
| Gib_CF2 | pCF203 | ATAATCTCCTCTATTTCCATGAGAATCACTCCTATTTTTTTG ATATATACATCATAACATTAC | This study |
| Gib_CF3 | pCF203 | AAAAAATAGGAGTGATTCTCATGGAAATAGAGGAGATTATAG AGAAAGTTGCTAGG | This study |
| Gib_CF4 | pCF203 | TGCGGGTCGTGGGGCGGGCGTCATCTCCCAAGCATTTTATGA GCCCTAGC | This study |
| Gib_CF5 | pSB1 | GCCGGTGGTTACCGTGATATTATCTATTACTATATCCCTATA TAAAGAATACTCAAAAAATGGGC | This study |
| Gib_CF6 | pSB1 | GTAATAGATAATATCACGGTAACCACCGGCTAGCAGGTGATG CATATGGCTAAAATGAGAATATCAC | This study |
| Gib_CF7 | pSV1_1 | TATTTTGAATCCATTGCGTTGCGCTCACTG | This study |

| | | | |
|---|---|---|---|
| Gib_CF8 | pSV1_1 | TGGGCGGCCGGCCGCGTTAATATTTTGTTAAAATTCGCGTTAAATTTTTGTTAAATCAG | This study |
| Gib_CF9 | pSV1_1 | AAATATTAACGCGGCCGGCCGCCC | This study |
| Gib_CF10 | pSV1_1 | TGCATTTTTTTGCGGCGCGCCCTGACA | This study |
| Gib_CF11 | pSV1_1 | AGCGCAACGCAATGGATTCAAAATAGATTCATAATGGAGTCATCCACG | This study |
| Gib_CF12 | pSV1_1 | TCAGGGCGCGCCGCAAAAAAATGCAAATAAAATTTGGGGTGG | This study |
| Res_CF1 | pSV1_2 | CGTACTGCAGCGATCGCGGTCATATGGATACAGCGGCC | This study |
| Res_CF2 | pSV1_2 | GTTATGGATTATAAGCGGCCGGC | This study |
| Gib_CF13 | pMVS1111A:$P_{synth}$-*bgaB* | CCACCCTGCCACCCCAAATTTTATTTGCATTTTTTTGCGGGTTAATTAAGCCTGGAGGAATGCCTTTATATAGG | This study |
| Gib_CF14 | pMVS1111A:$P_{synth}$-*bgaB* | TTTATATATTTTTAATTCACTGGGGGCAATTCTGTCAGGGCGCGCCTGGGGTCGTGCGCTC | This study |
| Gib_CF15 | pMVS1111A:$P_{hmtB}$/$P_{mrt(\mathit{M.t.})}$-*bgaB* | CGGCTCTAGCTATGTCCGATC | This study |
| Gib_CF16 | pMVS1111A:$P_{hmtB}$/$P_{mrt(\mathit{M.t.})}$-*bgaB* | CACTGGGGGCAATTCTGTCAG | This study |
| Gib_CF21 | pCF201 | AATACAAGAAAGGCGCGCCAAATCATTATATAGGACCTTGATAAAATTTTTTAGAGGGC | This study |
| Gib_CF22 | pCF201 | ACCTGACGTGTGGCCGGCCCGATTCAAATATAACAGCCGTTATAACACCGC | This study |
| Gib_CF23 | pCF201 | AATCGGGCCGGCCACACGTCAGGTGGCACTTTTCG | This study |
| Gib_CF24 | pCF201 | CATCCACGGATGCGATCGCCTAAGAAACCATTATTATCATGACATTAACCTATAAAAATAGGC | This study |
| Gib_CF17 | pCF202 | AAAAAATAGGAGTGATTCTCATGGCTAAAATGAGAATATCACCGGAAT | This study |
| Gib_CF18 | pCF202 | TGCGGGTCGTGGGGCGGGCGCTAAAACAATTCATCCAGTAAAATATAATATTTTATTTTCTCCCAAT | This study |
| Gib_CF19 | pCF202 | GATATTCTCATTTTAGCCATGAGAATCACTCCTATTTTTTTGATATATACATCATAACATTAC | This study |
| Gib_CF20 | pCF202 | TACTGGATGAATTGTTTTAGCGCCCGCCCCACG | This study |
| Res_LM1 | pMVS1111A:$P_{\mathit{hmtB}}$-$\mathit{fdh}_{\mathit{Z\text{-}245}}$ | ATCGGCTTAGGCGCGCCTGCTCATCGTCAATTCTAGTAGAGTCATGAATCATTATGCAGG | This study |
| Res_LM2 | pMVS1111A:$P_{\mathit{hmtB}}$-$\mathit{fdh}_{\mathit{Z\text{-}245}}$ | GCTTAGCGCATTAATTAACCGCCCATTTTTTGAGTATTC | This study |
| Gib_LM1 | pLM201 | TAACAGCGGCGCTATCAAGGTCCTGCATAATGATTCATGACGCCCGCCCCACG | This study |
| Gib_LM2 | pLM201 | TTTGCCGTATCTGCAGGCGATTTAAAAGATGATCCCATGAGAATCACTCCTATTTTTTTG | This study |
| Gib_LM3 | pLM201 | TTATGATGTATATATCAAAAAAATAGGAGTGATTCTCATGGGATCATCTTTTAAATCGCC | This study |
| Gib_LM4 | pLM201 | TCCTTTCGGTCGGGCGCTGCGGGTCGTGGGGCGGGCGTCATGAATCATTATGCAGGACC | This study |
| Gib_LM5 | pLM202 | ATCCTATATAAATATATCGCTAATTTTAAGGTTTTTCTGAGCCATCGGTTGGTTCATGGGTTAATTAAGAATACTCAAAAAATGGGCG | This study |
| Gib_LM6 | pLM202 | CGATATATTTATATAGGATTATATGAATAGATAATATCACATAAAATGAGGTGGTTAATTATGGGATCATCTTTTAAATCGCCTGCAG | This study |
| Seq_CF1 | specific for gDNA *M. t.*1.5 kb | CCACCAGTTCGACTCCCTGG | This study |
| Seq_CF2 | specific for gDNA *M. t.*1.5 kb | CTGTTAAAGGCGGGGGTGG | This study |
| Seq_CF3 | specific for gDNA *M. t.* 2.8 kb | CTTGGGTGATGATGGGATGTATTG | This study |

| | | | |
|---|---|---|---|
| Seq_CF4 | specific for gDNA *M. t.* 2.8 kb | CGAGGAGAAACACATCCAGCTG | This study |
| Seq_CF5 | specific for pME2001 replicon | GTTAATCCAGCACATCCTCC | This study |
| Seq_CF6 | specific for pME2001 replicon | CCTGTCCAACTTATACCTTTGG | This study |
| Seq_CF7 | analysis of *bgaB* constructs | CCCCATAACATCGGCACAGTAC | This study |
| Seq_CF8 | analysis of *bgaB* constructs | CCTGGCTGGGGTTAATAAATGTTG | This study |
| Seq_LM1 | analysis of *fdh*$_{Z\text{-}245}$ constructs | GATTTCTGGAATCCGCCATGGG | This study |
| Seq_LM2 | analysis of *fdh*$_{Z\text{-}245}$ constructs | CTAATAGTCGCCGATCCAAG | This study |
| Seq_LM3 | analysis of *fdh*$_{Z\text{-}245}$ constructs | GGTTCCTGGCTTGAATG | This study |
| Seq_LM4 | analysis of *fdh*$_{Z\text{-}245}$ constructs | GAGAAGCAAAGGATGACTG | This study |
| Seq_LM5 | analysis of *fdh*$_{Z\text{-}245}$ constructs | CAGCACCCATCTTATTCG | This study |
| Seq_LM6 | analysis of *fdh*$_{Z\text{-}245}$ constructs | GCAGTTAAGAAGGGTTCG | This study |
| Seq_LM7 | analysis of *fdh*$_{Z\text{-}245}$ constructs | GGCTCCGTTATAAGGGTTG | This study |
| Seq_LM8 | analysis of *fdh*$_{Z\text{-}245}$ constructs | CTGAATGGATCGAGAAAGG | This study |
| Seq_LM9 | analysis of *fdh*$_{Z\text{-}245}$ constructs | CATTCTTTCGAGATGGAAG | This study |
| Seq_LM10 | analysis of *fdh*$_{Z\text{-}245}$ constructs | CCTATATTCGCATTCGTGG | This study |
| Seq_LM11 | analysis of *fdh*$_{Z\text{-}245}$ constructs | ATGTTTGCCACACTGTG | This study |
| Seq_LM12 | analysis of *fdh*$_{Z\text{-}245}$ constructs | GGTGGGGTTTTGGTGTGCG | This study |

**Table S3.** List of gBlocks used in this study

| Name | Sequence (5' -> 3') | Reference |
|---|---|---|
| gBlock P$_{mcrB}$-*pac*-T$_{mcr}$ | GGTACCGAAAAGTGCCACCTGACCGATGGCCGGCCGCCCATTTTTTGAGTATTCAAATT<br>CAAATTATTGTGTTATTAACATCTTATATATAAACTTTTCTATTTAATGTTAATGAAAA<br>AGTGAATATATATACATAGAGTAATGTTATGATGTATATATCAAAAAAATAGGAGTGAT<br>TCTCATGACCGAGTACAAGCCCACCGTTAGGCTCGCAACCAGGGATGATGTTCCCAGGG<br>CAGTTAGGACCCTCGCAGCAGCATTCGCAGATTACCCCGCAACCAGGCACACCGTTGAT<br>CCCGATAGGCACATAGAGAGGGTTACCGAGCTCCAGGAGCTCTTCCTCACCAGGGTTGG<br>TCTCGATATAGGTAAGGTTTGGGTTGCAGATGATGGTGCAGCAGTTGCAGTTTGGACCA<br>CCCCCGAGTCAGTTGAGGCAGGTGCAGTTTTCGCAGAGATAGGTCCCAGGATGGCAGAG<br>CTCTCAGGTTCAAGGCTCGCAGCACAGCAGCAGATGGAGGGTCTCCTCGCACCCCACAG<br>GCCCAAGGAGCCCGCATGGTTCCTCGCAACCGTTGGTGTTTCACCCGATCACCAGGGTA<br>AGGGTCTCGGTTCAGCAGTTGTTCTCCCCGGTGTTGAGGCAGCAGAGAGGGCAGGTGTT<br>CCCGCATTCCTCGAGACCTCAGCACCCAGGAACCTCCCCTTCTACGAGAGGCTCGGTTT | This study |

| | | |
|---|---|---|
| | CACCGTTACCGCAGATGTTGAGTGCCCCAAGGATAGGGCAACCTGGTGCATGACCAGGA<br>AGCCCGGTGCATGACGCCCGCCCCACGACCCGCAGCGCCCGACCGAAAGGAGCGCACGA<br>CCCCATGGCTCCGACCGAAGCCACCCGGGGCGGCCCCGCCGACCCCGCACCCGCCCCCG<br>AGGCCCACCGCGGGGGACACACCGAACACGCCGACCCTGCTGAACACGCGGCGCAGTTC<br>GGTGCCCAGGAGCGGATCGGGAATTAATTCGAAGCTGCTGGTGAAAGAGACCCTATCTT<br>ACCTGCTAAAATCTAAGTTAATTACTAATTTATTATTAATTTATTATTAGATTGGGCAA<br>AATAGTAAAAGAAAACTAAAGGAAACCTAATATGGTTTCCTTTTTTTATATATTTTTAA<br>TTCACTGGGGGCAATTCTGTCAGGGCGCGCCTTCGGGCCATCGGGCCC | |
| gBlock $P_{hmtB}$-*Pac*I-*bgaB* | CGGCTCTAGCTATGTCCGATCAATCTTAATTAAGCCTGGAGGAATGCCCCCATGAACCA<br>ACCGATGGCTCAGAAAAACCTTAAAATTAGCGATATATTTATATAGGATTATATGAATA<br>GATAATATCACATAAAATGAGGTGGTTAATTATGAACGTTCTCAGTTCCATCTGCTATG<br>GGGGGGATTACAAC | This study |
| gBlock $P_{mrt(M.t.)}$-*Pac*I-*bgaB*-cor | CGGCTCTAGCTATGTCCGATCAATCTTAATTAAGCCTGGAGGAATGCCCCATTTCCATG<br>GATTATCGCTGGCAATCCCATAACCCCATCAGTTTTATTAATAAAATAGTAAATTTTAT<br>TAATAAATAAATAAAACAAGAGGTGTGAATACCATGAACGTTCTCAGTTCCATCTGCTA<br>TGGGGGGGATTACAAC | This study |
| gBlock codon optimized *bgaB* | CTGACAGAATTGCCCCCAGTGAATTAAAAATATATAAAAAAAGGAAACCATATTAGGTT<br>TCCTTTAGTTTTCTTTTACTATTTTGCCCAATCTAATAATAAATTAATAATAAATTAGT<br>AATTAACTTAGATTTTAGCAGGTAAGTGGGGTCGTGCGCTCCTTTCGGTCGGGCGCTGC<br>GGGTCGTGGGGCGGGCGCTAGACCTTGCCGGCTTCGTCGTGTTCCCTAAGGACAGCGAC<br>GTCGACGCCCTGAATCCTGAGTTCACCCCCCCTGAAGCATTTGCCATCTATCATATTCT<br>GGTAGATCTTATCTTCCGGAAGGGAGAGTGTGACCTCATAGTCGTTGTGGTTAATTATA<br>ATAAGGTACTTCCATTCATCGGTCTCCCTCTGCTGAACTTCGACATTCTCAGCAACCTC<br>CAGTATAGGATTTATGTGGTGTTTAGCAAACACCTGTTCGAGAAGCCTGCCAAGGTAGT<br>TGCTGTCAGGGTATGTTCCTACGTATATGCCCTCCCCCTTTCCGTAGCAGTTCCTGGTA<br>ACAGCAGGAAGGCCGGCATACCAATCACCTTTGAATGTGGCGAGAGGCTCAGCACCTTC<br>CAGCCTTATTATATCGGCCCATGTGGTACAGTCATACTCGCCGTCGTTTGAGTAGATCT<br>TATTCACCTTTGTCTCGGGATAAGGAACGAATTCCTCCACGAAGATGCCGAGAATGTCC<br>CTCAGCGGTCCTGGATATCCCCCGAGGTGCACTCTATCGTTCTCATCGACTATCACACT<br>GAAAAAGCTTACAATCAGGGTTCCGCCGTTTGCGACAAACTGCCTAAGGTTCTCATCTT<br>CTCCCTCTTTCACCATATACAGCATCGGTGCAATAACAACCTTATATTTTGTGAGATCG<br>TCGGACGGTCTTACAAAGTCGACTGCTATGTTTCTCTTGTAAAGCTCTCTATAATATGC<br>CTCTACTATGGGAATATATCTGAGCTTGTTGTGCGGTTTGGAACTGAGCTCAACTGCCC<br>ACCAGTTTTCCCAGTCAAAGATAATTGCCACCTCTGCCTTTATTCTACTCCCCACGAGG<br>CAGTCAAGTTTTTTCAGCTCCTGGCCAAGCTGGGTAACTTCCCTGTATATTCTATTGTT<br>TTCGTTAAGAAAGTGGGGCACCATTGCTCCGTGAAACTTCTCAGCTCCTGCTCTGGACT<br>GCCTCCACTGAAAGAACATTATCCCATCGGCACCCCTGGCGATTGTTGCGTAACTCCAG<br>AGTCTCATAACCCCCGGCGGCTTTGGCACATTGATATCTCTCCAATTAACGTGACTGGT<br>GACCTGCTCCATAAGAATGAACGGCTGCCCCTTCCTAAGTGACCTCATGAGGTCATTCA<br>TCATTGCGTGCTGTATAGGGAGTCCCTCCCTGGGATCTGGGTAGCTATCCCAGGTAACG<br>ATATCTACGTGCTGAGCCCACTGAAAGTAGTTGAGTGGCTTGAATGATCCCATGAAATT<br>TGTGGAGACCGGGATATCGGGGGTTACTTCCCTGAGGATCTCCTTTTCTGTAAGGAAGA<br>GTTTGAGGATTGAATCATTCATGAATCTGTAGTAATCAAGCTCCTGGCTGGGGTTAATA<br>AATGTTGGTGCCTTCCTAGGGGGATTAATCTCATCCCAGTGGTTATATCTCTGGCCCCA | This study |

GAAGTTTGTACCCCATCTTTCATTAAGTTCATCAATGGTCTTATACCTTTCTTTAAGCC
ATTTTCTGAAAGCAACTGCGCAATTCTCACAGAAACACTTACTTACATGGCAAGCGTAT
TCGTTATTTACGTGCCACATTTTGAGGGCTGGATGATTTTTGTATCTCTCAGCTATAGC
CCTTACCAGCCTCTTTATATGTGTTATAAGCTGAGGGTGATTTGGGCAATAATGCTGTC
TACTCCCGAAACTCAGTATCACACCGGACTCGTCAATAGGGAGTGAATCAGGGTATTTC
TTCACGAACCAGGCGGGTGTGGTTGCGGTGGCGGTCCCCAGATTTATGTATACCCCATG
ATCGTAGAGGATGTCTATCACTTTGTCGAGCCATTCAAAATCAAATACACCGTCTGATG
GCTCGATTTTGGACCAGCTAAAGATTCCGAGTGAAACAAGATTAACACCGGCCTTCTGC
ATAAGTTTTGCGTCCTCGTACCATATCTCCTCGGGCCACTGTTCTGGGTTGTAATCCCC
CCCATAGCAGATGGAACTGAGAACGTTCATATGCATCACCTGCTAGCCGGTGGTTACCG
TGATATTATCTATTACTATATCCCTATATAAAGGCATTCCTCCAGGCTTAATTAAC

**Table S4.** List of plasmids used in this study

| **Name** | **Function** | **Reference** | ***M. t.* strain** |
|---|---|---|---|
| pMTL83151 | Shuttle vector for *Clostridia* spp. | Heap *et al.* (2009) | - |
| pMU131 | Shuttle vector for *Thermoanaerobacter* spp. | Shaw *et al.* (2010) | - |
| pME2001 | Cryptic plasmid of *M. marburgensis* | Bokranz *et al.* (1990) | - |
| pBBR1-MCS2 | Standard cloning vector in *E. coli* | Kovach *et al.* (1995) | - |
| pUC19 | Standard cloning vector in *E. coli* | Yanisch-Perron *et al.* (1985) | - |
| pYS3 | Shuttle vector for *Pyrococcus furiosus* including Sim$^r$ | Waege *et al.* (2010) | - |
| pME2508 | PeiP production in *E. coli* | Luo *et al.* (2002) | - |
| pCF200 | pUC57 vector including synthesized $P_{mcrB(M.v.)}$_Pur$^r$ codon-optimized for *M. thermautotrophicus*, $T_{mcr}$ | This study | - |
| pCF201 | pUC19 vector including native *M. thermautotrophicus* Z-245 *fdh*$_{Z\text{-}245}$ operon with putative promoter region | This study | - |
| pLM201 | Exchange of Neo$^r$ to coding region of *fdh*$_{Z\text{-}245}$ from pCF201 in pCF204 | This study | - |
| pLM202 | Exchange of $P_{mcrB(M.v.)}$ to $P_{hmtB}$ in pLM201 | This study | - |
| pCF203 | Exchange of Pur$^r$ to Sim$^r$ in pCF200 | This study | - |
| pCF204 | Exchange of Pur$^r$ to Neo$^r$ in pCF200 | This study | - |
| pCF404 | pUC57 including 1 kb up- and downstream of annotated *pyrF* gene (MTH_RS00570) and $P_{mcrB(M.v.)}$_Pur$^r$ | This study | - |
| pCF407 | Exchange of $P_{mcrB(M.v.)}$_Pur$^r$ to $P_{mcrB(M.v.)}$_Neo$^r$ in pCF404 | This study | - |

| | | | |
|---|---|---|---|
| pSB1 | Exchange of $P_{mcrB(M.v.)}$ promoter to $P_{synth}$ in pCF407 | This study | - |
| pSV1_1 | Shuttle vector construct containing $P_{mcrB(M.v.)}$_Sim$^r$ and pBBR1MCS2 backbone and pME2001 replicon | This study | - |
| pSV1_2 | Shuttle vector construct containing $P_{mcrB(M.v.)}$_Sim$^r$ and pMTL backbone and pME2001 replicon | This study | - |
| pSV1_3 | Shuttle vector construct containing $P_{mcrB(M.v.)}$_Neo$^r$ and pMTL80151 backbone and pME2001 replicon | This study | - |
| pMVS-V1 | Shuttle vector construct containing $P_{synth}$_Neo$^r$ and pMTL backbone and pME2001 replicon | This study | x |
| pMVS1111A: $P_{synth}$-*bgaB* | Shuttle vector construct pMVS-V1 including β-galactosidase (*bgaB*)-gene and promoter $P_{synth}$ | This study | x |
| pMVS1111A: $P_{hmtB}$-*bgaB* | Shuttle vector construct pMVS-V1 including β-galactosidase (*bgaB*)-gene and promoter $P_{hmtB}$ | This study | x |
| pMVS1111A: $P_{mrt(M.t.)}$-*bgaB* | Shuttle vector construct pMVS-V1 including β-galactosidase (*bgaB*)-gene and promoter $P_{mrt(M.t.)}$ | This study | x |
| pMVS1111A: $P_{synth(BRE)}$-*bgaB* | Shuttle vector construct pMVS-V1 including β-galactosidase (*bgaB*)-gene and promoter $P_{synth(BRE)}$ | This study | x |
| pMVS1111A: $P_{hmtB}$-$fdh_{Z\text{-}245}$ | Shuttle vector construct pMVS-V1 including $fdh_{Z\text{-}245}$ operon from *M. thermautotrophicus* Z-245 and promoter $P_{hmtB}$ | This study | x |

### Cultivation

*Methanothermobacter* spp. were cultivated according to basic principles for methanogen cultivation as stated in Balch *et al.* (1979), with adjustments to state-of-the-art anaerobic handling equipment. The mineral medium contained (per liter): sodium chloride, 0.45 g; sodium hydrogen carbonate, 6.00 g; di-potassium hydrogen phosphate, 0.17 g; potassium di-hydrogen phosphate, 0.23; ammonium chloride, 0.19 g; magnesium chloride, 0.08 g; calcium chloride dihydrate, 0.06 g; ammonium nickel sulfate, 1 mL (0.2 weight%); iron(II)chloride pentahydrate, 1 ml (0.2 weight%); resazurin indicator solution, 4 mL (0.025 weight%); and trace element solution, 1 mL (10-fold as stated in Balch and Wolfe (1976)). All chemicals were *p.a.* grade. No vitamin solution was added. For the formate growth experiments, the mineral medium was supplemented with 100 mM sodium formate, 10 µM sodium molybdate, 1 µM sodium selenite, and 0.125 weight% yeast extract. The medium was gassed using $N_2/CO_2$ (80/20 volume%) to eliminate dissolved oxygen. The pH value was adjusted to 7.2 using hydrochloric acid. As reducing agent and sulfur source 0.5 g/L cysteine hydrochloride, and for solid mineral medium additionally 0.3 g/L sodium sulfide monohydrate, was added. Afterwards, the mineral medium was dispensed into serum bottles inside of an anaerobic chamber with a 100% $N_2$ atmosphere (UniLab Pro Eco, MBraun, Garching, Germany). The headspace of the serum bottles was exchanged to 200 kPa $H_2/CO_2$ (80/20 volume%) and autoclaved (100 kPa, 20 min, 121°C). For the formate growth experiments, the headspace of the serum bottles was exchanged to 152 kPa $H_2/CO_2$ (80/20 volume%), which provides the same electron equivalents compared to the 100 mM formate as positive control, or 152 kPa $N_2/CO_2$ (80/20 volume%). All *Methanothermobacter* strains in liquid medium were incubated at 60°C with shaking at 150 rpm (Lab companion ISS-7100R, Jeio Tech, Republic of Korea). For cultivation of genetically modified *M. thermautotrophicus* ΔH strains, 250 µg/mL neomycin sodium salt was added. For solidified mineral medium, 1.5 weight% Bacto™ agar (BD Life Science, Berkshire, UK) was supplemented *prior* to autoclaving. Afterwards, solidified media plates were poured and dried for two hours inside of the anaerobic chamber. *M. thermautotrophicus* ΔH was applied to solidified media plates by spot-plating, spread-plating, or pour-plating. For spot-plating, 50 µL of *M. thermautotrophicus* ΔH culture was spotted on a solidified media plate. Incubation was started after the drop was completely absorbed. For spread-plating, 50 µL of diluted or undiluted liquid culture was applied to a solidified media plate and spread out with a Drigalski spatula until the liquid was completely absorbed. For pour-plating, 5 mL mineral medium containing 0.8 weight% Bacto agar (soft-agar) were mixed with a liquid *M. thermautotrophicus* ΔH culture and poured on top of a solidified media plate, which contained 1.5 weight% Bacto agar. For better gas-solid mass transfer and to avoid sealing of the plates by water, paper clips were added to the petri dish *prior* to incubation in a custom-made stainless-steel jar (Raff + Grund, Freiburg am Neckar, Germany) inspired by Balch *et al.* (1979). The gas phase of the stainless-steel jar was exchanged to 200 kPa of $H_2/CO_2/H_2S$ (79.9/20/0.1 volume%). The pressurized anaerobic jar was incubated without shaking at 60°C (Memmert, Schwabach, Germany).

For general cloning and gene manipulation, *E. coli* NEB stable (New England Biolabs, Frankfurt/Main, Germany) was used. *E. coli* S17-1 for conjugational DNA transfer was kindly provided by Prof. Dr. Wolfgang Wohlleben of the Department for Biotechnology at the University of Tübingen, Germany (25). *E. coli* BL21(DE3) with pME2508 (Archaea Center of the University of Regensburg, Germany) was used to produce recombinant PeiP enzyme. *E. coli* was cultivated in LB medium, which contained (per liter): sodium chloride, 10 g; trypone, 10 g; yeast extract, 5 g, and which was supplemented with appropriate amounts of chloramphenicol (30 µg/mL), ampicillin (100 µg/mL), or kanamycin (50 µg/mL). For cultivation of *E. coli* S17-1, trimethoprim (10 µg/mL) was added to stabilize the genome-integrated *tra* module, which is responsible for mobilization of plasmid DNA (25). Solidified LB media plates contained 1.5 weight% of Kobe I Agar (Carl Roth, Karlsruhe, Germany) and were incubated at 37°C. Liquid *E. coli* cultures were incubated at 37°C with shaking at 150 rpm.

**Molecular methods and construction of modular shuttle-vector system (pMVS)**

PCR was performed with Q5 Hot Start High fidelity polymerase (NEB, Ipswitch MA, USA) according to the manufacturer's guidelines and with the required primer combinations (**Table S2**). Primer concentrations were reduced 10-fold and elongation time was prolonged by 1 min. Resulting PCR products were *Dpn*I digested when required, and purified using a PCR purification Kit (Qiagen, Hilden, Germany). For initial fusion of the first shuttle-vector construct pSV1_1, as described below, we used the Gibson Assembly® Ultra kit (Synthetic genomics, La Jolla CA, USA). All follow-up constructs were assembled with Gibson® Assembly Master Mix (NEB, Ipswitch MA, USA) or restriction/ligation cloning with the aid of the implemented modular restriction enzyme-recognition sites (**Figure 1**). *E. coli* cells were transformed with DNA *via* chemical transformation by following a standard heat-shock protocol (26). All plasmids were confirmed by Sanger sequencing (MPI genomics center, Tübingen, Germany).

pCF200, which contains the puromycin acetyltransferase (*pac*)-gene ($Pur^r$) from *Streptomyces alboniger* as a codon-optimized version for *M. thermautotrophicus* ΔH under the control of the $P_{mcrB(M.v.)}$ promoter and the $T_{mcr}$ terminator from *Methanococcus voltae* (1), was completely synthesized (BioCat, Heidelberg, Germany). The *pac*-gene in pCF200 was exchanged to the 3-hydroxy-3-methylglutaryl-coenzyme A reductase (HmgA)-encoding gene ($Sim^r$) from *Thermococcus kodakarensis* by using pYS3 (21) and pCF200 as templates for Gibson® Assembly, resulting in pCF203. pCF203, pME2001 (extracted from wild-type *M. marburgensis*), and pBBR1-MCS2 (Addgene #85168) were used as templates for Gibson® Assembly with the Gibson Assembly® Ultra kit (Synthetic genomics, La Jolla CA, USA), and resulted in the putative shuttle vector pSV1_1. pSV1_1 was the basis for further shuttle vectors. For the introduction of a high copy number replicon for *E.*

*coli,* a *tra*-region for plasmid mobilization, and an additional *AsiS*I restriction enzyme-recognition sequence, the pBBR1-MCS2 backbone was exchanged to the *E. coli* vector backbone from pMTL83151 (5), including Cam$^{r}$, ColE1, and the *tra* minigene for mobilization, *via* Gibson® Assembly resulting in pSV1_2. To implement the thermostable neomycin phosphotransferase gene (Neo$^{r}$) (27), the Pur$^{r}$ from pCF200 was exchanged to Neo$^{r}$ from pMU131 (10) by Gibson® Assembly. Afterwards, based on pCF404, the Neo$^{r}$ under the control of the $P_{\mathit{mcrB(M.v.)}}$ promoter and the $T_{\mathit{mcr}}$ terminator, was used to construct a putative integration plasmid for the exchange of an annotated *pyrF* gene in *M. thermautotrophicus* ΔH, using *Asc*I and *Fse*I as restriction enzymes and T4 ligase for ligation, resulting in pCF407. In pCF407 the $P_{\mathit{mcrB(M.v.)}}$ promoter was exchanged to $P_{\mathrm{synth}}$ by inverse PCR resulting in pSB1. The fragments $P_{\mathit{mcrB}}$_Neo$^{r}$_$T_{\mathit{mcr}}$ from pCF407 and $P_{\mathrm{synth}}$_Neo$^{r}$_$T_{\mathit{mcr}}$ from pSB1 were used to substitute the Sim$^{r}$ in pSV1_2 by restriction-ligation cloning using *Asc*I and *Fse*I, resulting in pSV1_3 and pMVS-V1, respectively. To generate pMVS1111A:$P_{\mathrm{synth}}$-*bgaB*, the PCR amplified gBlock with the thermostable β-galactosidase (*bgaB*) gene, which was codon-optimized for *M. thermautotrophicus* ΔH and which was placed under the control of the $P_{\mathrm{synth}}$ promoter, was fused to *Asc*I-digested pMVS-V1 with Gibson® Assembly. The *Asc*I restriction enzyme-recognition sequence was recovered at the intersection with the selectable-marker module and a *Pac*I sequence was introduced at the intersection with the *M. thermautotrophicus* ΔH-replicon module. Further promoters ($P_{\mathrm{synth(BRE)}}$, $P_{\mathit{hmtB}}$, $P_{\mathit{mrt(M.t.)}}$) were amplified *via* overlap-extension PCR of the β-galactosidase gBlock and promoter gBlock and inserted by restriction/ligation cloning using restriction enzymes *Pac*I and *Asc*I. pCF201 was constructed by amplifying the $\mathit{fdh}_{\mathrm{Z\text{-}245}}$-operon from *M. thermautotrophicus* Z-245 genomic DNA and introducing the fragment into pUC19 by Gibson® Assembly. Gibson® Assembly was used to exchange the Neo$^{r}$-coding region in pCF204 with the $\mathit{fdh}_{\mathrm{Z\text{-}245}}$-operon from pCF201, resulting in plasmid pLM201. The promoter $P_{\mathit{mcrB(M.v.)}}$ in pLM201 was exchanged to $P_{\mathit{hmtB}}$ by inverse PCR of the complete plasmid, except of the $P_{\mathit{mcrB(M.v.)}}$ sequence, with primers containing overlapping parts of $P_{\mathit{hmtB}}$ in the overhangs, and direct transformation of *E. coli* with the linear PCR product. The resulting plasmid pLM202 was used to amplify the $P_{\mathit{hmtB}}$-$\mathit{fdh}_{\mathrm{Z\text{-}245}}$ cassette by PCR to include *Pac*I and *Asc*I restriction enzyme-recognition sequences. Restriction/ligation cloning with the restriction enzymes *Pac*I and *Asc*I was used to exchange the β-galactosidase gene in pMVS1111A:$P_{\mathrm{synth}}$-*bgaB* for the $P_{\mathit{hmtB}}$-$\mathit{fdh}_{\mathit{Z\text{-}245}}$ cassette to give pMVS1111A:$P_{\mathit{hmtB}}$-$\mathit{fdh}_{\mathit{Z\text{-}245}}$.

**Plasmid-DNA extraction from *Methanothermobacter* spp.**

For plasmid-DNA extraction from *Methanothermobacter* spp., 10 mL of liquid cell culture was centrifuged at 3700 rpm for 15 min at room temperature (Centrifuge 5920 R, rotor S-4x1000, Eppendorf, Hamburg, Germany). The supernatant was discarded and the cell pellet was resuspended in 150 µL of sucrose (30 weight%)-containing Buffer P1 (from QIAprep Spin Miniprep Kit, Qiagen, Hilden, Germany). For lysis of *Methanothermobacter spp.* cells, alkaline lysis was combined with enzymatic lysis by adding a final concentration of 100 ng/mL of pseudomurein endoisopeptidase (PeiP) to the sample *prior* to incubation for 1 h at 60°C. The pseudomurein-degrading enzyme PeiP, which lyses pseudomurein-containing Methanobacteriales cell walls, was produced as a heterologous 6xHistidine-tagged protein from pME2508-carrying *E. coli* BL21(DE3) as described (22). The recombinant protein was purified *via* a Protino Ni-TED column according to the manufacturer's guidelines (Macherey+Nagel, Düren, Germany). After the PeiP treatment, the QIAprep Spin Miniprep Kit (Qiagen, Hilden, Germany) manufacturer's guidelines were followed with final elution in 40 µL nuclease-free water.

**Interdomain conjugational DNA transfer**

*E. coli* S17-1 was transformed with the respective shuttle vector. Over-night cultures of the respective *E. coli* S17-1 donor strains were inoculated. At the same time, 20 mL of liquid mineral medium was inoculated with wild-type *M. thermautotrophicus* ΔH (recipient). The over-night culture of *E. coli* S17-1, which contained the shuttle vector, was diluted into 10 mL of fresh LB medium in a sterile 50-mL baffled flask for better aeration to give an $OD_{600}$ of 0.3-0.5. When this culture reached an $OD_{600}$ of 2.0-2.5, the incubation was stopped and the culture was harvested aerobically at 3700 rpm for 10 min at room temperature (Centrifuge 5920 R, rotor S-4x1000, Eppendorf, Hamburg, Germany). The supernatant was discarded and the *E. coli* S17-1 pellet was transferred into the anaerobic chamber. Wild-type *M. thermautotrophicus* ΔH was grown to early stationary growth phase ($OD_{600}$ of 0.25-0.35). 8 mL of *M. thermautotrophicus* ΔH culture was centrifuged stepwise at 12500 rpm for 4 min at room temperature (Centrifuge 5424, rotor FA-45-24-11, Eppendorf, Hamburg, Germany) inside the anaerobic chamber. The final pellet was resuspended in 250 µL of the original non-concentrated *M. thermautotrophicus* ΔH culture, and gently mixed with the *E. coli* S17-1 pellet. 100 µl of cell suspension were anaerobically spotted on solid LB-MS medium, which was a mixture that consisted of 50 volume% of mineral medium and 50 volume% of LB medium without the 10 g/L sodium chloride. The spot was dried, while the lid of the petri dish was kept slightly open for 1 h at 37°C in the incubator (Coy laboratory products, Green Lake MA, USA) within the anaerobic chamber. When the spot was completely absorbed, the plates were provided with paper clips and transferred to a stainless-steel jar. The gas phase of the jar was exchanged to 200 kPa $H_2/CO_2/H_2S$ (79.9/20/0.1 volume%) and incubated at 37°C without shaking for 16-20 h. The spot-mated *E. coli* S17-1 and *M. thermautotrophicus* ΔH cells were washed from the LB-MS plates using 1 mL non-selective mineral medium and transferred to 4 mL non-selective mineral medium in a 50-mL serum bottle with a $H_2/CO_2$ (80/20 volume%) gas phase. After recovery for 3-4 h at 60°C with shaking at 150 rpm, 1 mL of the culture was transferred to 20 mL selective liquid mineral medium in a 100-mL serum bottle and incubated at 60°C with shaking at 150 rpm. Growth of *M. thermautotrophicus* ΔH after 24-48 h of incubation indicated successful DNA transfer into *M. thermautotrophicus* ΔH, while growth only later than 48 h indicated the appearance of spontaneously neomycin-resistant *M. thermautotrophicus* ΔH cells. 50 µL from this selective-enrichment culture was spread-plated on selective solidified media plates, and individual colonies were analyzed after two days of incubation at 60°C.

To determine the conjugation frequency, the following modifications to the standard protocol were made: **1)** the cell count of *M. thermautotrophicus* ΔH in liquid culture was determined by counting in a Petroff-counting chamber. The initial recipient cell number in 100 µL of the stepwise-concentrated culture was calculated based on this cell count; **2)** the 5 mL non-selective recovery culture from the washed spot after spot-mating was incubated for ~16-20 h instead of 3-4 h; and **3)** 100 µL of the non-selective recovery culture was directly spread-plated on selective solidified media plates, without a liquid selective-enrichment step.

#### Molecular methods for analysis of genetically modified *M. thermautotrophicus* ΔH

PCR analysis was performed from liquid cultures and directly from individual colonies. 100 µL of liquid culture or one individual colony, which was resuspended in 40 µL of deionized water, were boiled for 12 min at 100°C (ThermoMixer C, Eppendorf, Hamburg, Germany). After cooling the sample on ice, 1 µL was added to a 10-µL PCR reaction mix. PCR was performed using Phire plant PCR master mix (Thermo

Scientific, Waltham MA, USA). The denaturation and annealing times were increased from 5 sec to 20 sec and to 10 sec, respectively. 30 cycles were performed for all analyses. We observed false positive PCR signals for shuttle-vector DNA due to plasmid DNA carry-over from *E. coli* for two transfers after the non-selective liquid recovery step. After the third transfer, plasmid DNA from *E. coli* was not detectable anymore in any of our experiments. For robust PCR amplifications of individual colonies from *M. thermautotrophicus* ΔH, it was crucial to keep the agar contamination of the PCR sample as little as possible. Therefore, even though the plating efficiency is higher with pour-plating, genetically modified *M. thermautotrophicus* ΔH strains were spread-plated instead of pour-plated. This led to a lower total colony count, but to more reliable results.

Additional to PCR analysis, plasmid DNA from genetically modified *M. thermautotrophicus* ΔH strains was extracted as described above. The purified plasmid DNA was used for retransformation of *E. coli* NEB stable. Analysis of *E. coli* NEB stable colonies was performed *via* test restriction digestions and Sanger sequencing for further confirmation of stable replication of shuttle vectors in *M. thermautotrophicus* ΔH.

**β-Galactosidase enzyme assays**

For a β-galactosidase enzyme activity assay with the lactose analogue S-Gal, 2 mL of over-night cell cultures that carry pMVS-V1 or pMVS1111A:$P_{Synth}$-*bgaB* were harvested by centrifugation for 4 min at 13000 rpm at room-temperature (Centrifuge 5424, rotor FA-45-24-11, Eppendorf, Hamburg, Germany). The supernatant was discarded and the samples were stored at -20°C until further use. All samples were resuspended in 100 µL Buffer P1 (from Qiagen QIAprep Spin Miniprep Kit) containing sucrose (30 weight%), and lysed by adding 100 ng/mL PeiP, followed by incubation for 30 min at 60°C. 50 µL of the cell lysate were incubated with 250 µg/mL S-Gal and 250 µg/mL ammonium ferric citrate in 1 mL LB medium, which provided any potentially required trace compounds. The samples were incubated for 1 hour at 60°C. After ~30 min, a color change was visible.

For a β-galactosidase enzyme activity assay with the lactose analogue ONPG, 4 mL of cell culture were harvested anaerobically by step-wise centrifugation (Centrifuge 5424, rotor FA-45-24-11, Eppendorf, Hamburg, Germany). Afterwards, the same lysis procedure for samples was applied as for the S-Gal assay. The resulting cell lysate was used for a quantitative *in-vitro* β-galactosidase enzyme activity assay with ONPG as chromogenic substance according to Jensen *et al.* (2017). In brief, 12.5 µL of cell lysate (equals to 0.5 mL of original cell culture) were mixed with 600 µL of ONPG (1 mg/mL)-containing substrate solution. The mixture was incubated for 2 h at 60°C. Afterwards, 200 µL were added to 200 µL of 1 M sodium bicarbonate stop-solution in a 96-well plate. The absorbance at 420 nm was measured in a micro-plate reader (Multiskan Go, Thermo Scientific Waltham MA, USA). For the preliminary experiment (**Figure S7**), 25 µL of cell lysate were mixed with 675 µL of ONPG substrate solution instead. After the incubation for 2 h at 60°C, 350 µL of substrate solution were added to 350 µL of stop solution, and the absorbance at 420 nm was measured in a cuvette (1-cm path length) with a spectrophotometer (NP80, Implen, Munich, Germany).
